# Crosslinked agarose-gelatine beads as a substrate for investigating biofilms of bacterial pathogens

**DOI:** 10.1101/2024.08.16.608334

**Authors:** Dan Roizman, Maren Herzog, Arpita Nath, Nivetha Pachaimuthu, Ahmad Hujeirat, Benno Kuropka, Jens Rolff, Alexandro Rodríguez-Rojas

## Abstract

Treating chronic bacterial infections is challenging due to the formation of biofilms, making bacteria less susceptible to antimicrobials. *In vitro* models have limitations in replicating biofilm physiology. To address this problem, we have created a hydrogel substrate that combines crosslinked agarose and gelatine presented as beads, providing stability and resistance to autoclaving. Bacterial pathogens rapidly colonise these biogel beads when submerged in liquid culture. The substrate was tested with *Escherichia coli*, *Pseudomonas aeruginosa*, and *Staphylococcus aureus*, showing more robust biofilm growth than its glass bead counterpart. Additionally, this led to increased virulence factor production and served as a reservoir for biofilm quorum sensing molecules. These features closely resemble clinical situations, suggesting a more accurate representation of biofilm-associated infections than current approaches. This new substrate offers a practical and convenient model for studying biofilms of bacterial pathogens, providing an efficient solution to the research community and holding promise for future breakthroughs.

## Introduction

Biofilms are microbial communities embedded in a self-produced extracellular gelatinous matrix. According to fossil records, they date back 3.5 billion years and facilitate cell adhesion and surface attachment [1,2]. Biofilms are ubiquitous and can be found in diverse habitats, ranging from the mammalian gut to deep subsurface rocks [3]. It is a predominant lifestyle on Earth, and estimations say 40-80% of bacterial and archaeal cells exist in biofilm form [4].

Bacterial biofilms are particularly associated with chronic infections and can withstand high antibiotic concentrations [5]. Biofilms are implicated in infections related to medical devices and other persistent conditions [6]. During infection, bacterial biofilms are often associated with treatment failure [7]. The lack of consensus among the diverse biofilm cultivation methods and analysis techniques further complicates the issue [8].

The most widely used method for biofilm formation involves microtiter plates; initially developed by Fletcher in 1977 for studying bacterial attachment, this technique has since been refined and used for quantifying biofilm biomass [9]. Several methodologies exist to study biofilm formation under simulated natural conditions. One example is the modified Robbins device [10], which utilises a flow system to circulate liquid-containing microorganisms [11]. Advances in fluorescence-based microscopy, such as confocal microscopy, have significantly contributed to our understanding of biofilm mechanisms when combined with later open [12] and closed flow chamber systems [11]. However, a significant limitation of contemporary systems is that they are conducted on artificial surfaces, primarily glass or plastic substrates [13]. Other materials such as silicone, Teflon, PTFE, polypropylene, titanium, or other metals such as stainless steel or copper are also commonly used. Data extrapolated from research on biofilm bactericidal compounds conducted on abiotic surfaces, such as polycarbonate filters, may not be entirely applicable to biofilms growing on biotic surfaces, such as skin or mucoid tissues, due to ultrastructural and physiological variations. These limitations underscore the need for new approaches to studying biofilms [14].

In 1881, Robert Koch used gelatine to develop the first solid media. However, its practicality was compromised due to its susceptibility to bacterial digestion and melting temperature below 37°C. These limitations prompted a search for alternative agents. A suitable substitute was found when Angelina Hesse, the wife and assistant of Walther Hesse, an associate of Robert Koch, proposed using agar to prepare solid cultures in 1887 [15]. Agar offered several advantages over gelatine, including resistance to bacterial degradation and a higher melting temperature, making it more practical for microbial growth [16]. Apart from its stability, agar alone is not an accurate model for recapturing conditions closer to *in vivo* infections [17]. Nonetheless, procedures based on fast planktonic growth and single colony isolation at 37°C rendered gelatine obsolete, although it represented the closest proxy to a host environment [16]. Notably, bacterial pathogens discovered and described in that era using Koch’s postulates still comprise most of the priority list 150 years later [18,19]. The increased challenges of handling those infections necessitate a paradigm shift. Its rationale is twofold: balancing the focus between biofilm and planktonic lifestyles and having better substrates that mimic organic surfaces, making them more comparable to within-host scenarios.

Agarose, a purified derivative of agar, is used to manufacture chromatography media for biomolecule purification through crosslinking its polysaccharide molecules [20,21]. The same crosslinking method serves as the basis for the work presented here. Our goal was to enhance the biocompatibility of the substrate by incorporating gelatine, inspired by an emulsification method to create bacteria-embedded agarose beads [22–24]. Additionally, we found advantages in the substrate shape of the *in vitro* biofilm models that employ glass beads as a substrate [25–27]. The ability of bacteria to attach to glass beads and grow, forming aggregates or biofilms, was well documented as early as the 1940s by researchers like Heukelekian and Heller in 1940 [28] and Zobell in 1943 [29], even before the term “biofilm” was coined. Building on previous research on biofilms, we develop a biological hydrogel-based substrate to enhance existing *in vitro* biofilm models for pathogenic bacteria. Our method combines and stabilises the advantages of agar and gelatine, the two classic substrates for bacterial growth in semisolid media. We also took advantage of the protocols established for glass bead biofilm [25] and accordingly shaped our substrate as a bead.

We chose three well-established bacterial models to study the biofilms of bacterial pathogens on the surface of crosslinked agarose gelatine. Specifically, we examined reference strains from three bacterial families of the ESKAPE group [30], consistently listed as major concern pathogens [18,31]. Among these are two notable pathogenic bacteria, one gram-negative and one gram-positive: *Pseudomonas aeruginosa* and *Staphylococcus aureus*. We also employed the non-pathogenic *Escherichia coli* K-12 model, the best-established bacterial model system since its initial isolation a hundred years ago [32]. Although *E. coli* is generally considered a moderate biofilm former, its extensive study history allows for in-depth analysis due to all genetic toolboxes and the accuracy of modern genome databases such as EcoCyc [33].

## Results

### Generation of agarose-gelatine beads and properties

First, we have focused on the two traditional substrates for semisolid media in microbiology: agarose and gelatine matrices. Due to their biological compatibility and low reactivity, we prepared a crosslinked polymer based on these materials to serve as a substrate. We explored various concentrations of reagents and the crosslinking conditions to create a set of conditions that could be used as a starting point. The method we adopted, based on the catalytic crosslinking activity of divinyl sulphone (DVS), which was first demonstrated by Porath, Låås & Janson [21], which, immediately following adaptation, showed great flexibility for diverse experiments in biofilm analysis. We attempted to develop optimal properties that would enable the fabrication of spherical beads. Combining 2% agarose and 2% gelatine produced the most favourable results for generating biogel spherical structures. Lower concentrations (ranging from 0.5 to 1%) led to easily deformable beads, while higher concentrations (ranging between 3 and 8% each) solidified rapidly, posing handling challenges. These structured, spherical beads were formed by depositing desired volumes of the hot mixture of agarose-gelatine in ice-cold mineral oil, as illustrated in Fig. 1A-E. Some preliminary experiments also confirmed that agarose-alone beads showed a lower biofilm growth than the combined beads. Gelatine-alone beads showed lower stiffness and deficient crosslinking compared to agarose or agarose-gelatine combination.

**Fig. 1.**
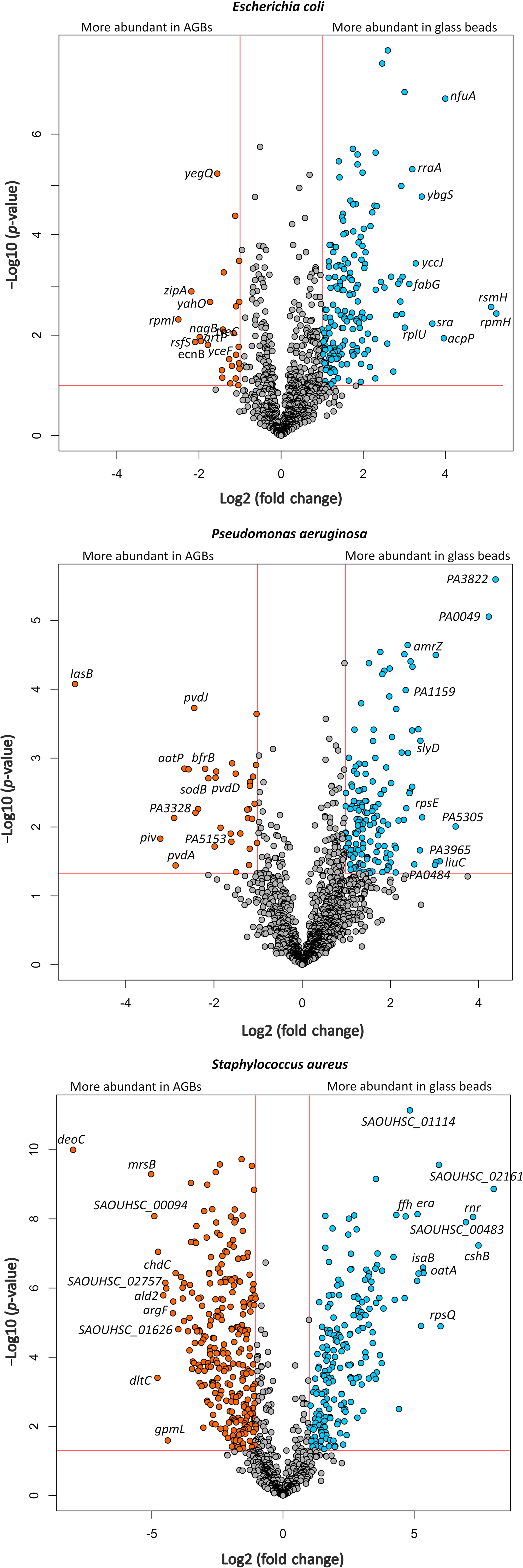
A setup for generating crosslinked agarose-gelatine beads and their properties. In the initial step (A), a mixture of gelatine-agarose (2% each in distilled water) is prepared and placed on a magnetic stirrer with a hot plate to maintain liquidity. A peristaltic pump continuously pumps this solution, forming drops deposited into ice-cold mineral oil to create millimetre-scale beads. Subsequently (B), the mineral oil is removed, and washing steps, including detergent, are performed to eliminate oil residues. Following this (C), the beads undergo crosslinking using divinyl sulphone (DVS), with the crosslinker later removed through additional washing steps. After removal of the crosslinker, the beads are transferred to a bottle and autoclaved at 121°C for 15 minutes at 1 atmosphere (D). Once autoclaved, the beads can be stored for several months until use. In the final step (E), beads are transferred to a 24-well multiwell plate, where the chosen bacteria are inoculated, and the biofilm is established, as previously described. Swelling stability post-crosslinking and autoclaving for five months in two standard culture media: LB (F) and MHB (G). The bars represent the mean of 5 beads (N=5), and the error bars represent the standard deviation. The crosslinking significantly increased the stiffness of the beads as measured via Young’s modulus of an atomic force microscope (H). We performed a student’s t-test, and the four asterix indicate that the p-values were below 0.0001. Comparative surface view of AGB and glass bead acquired with a Keyence VHX-X1 microscope (I). See the Material and Methods section and supplementary video S1 for additional details.

Initially, we used a multi-stepper pipette to dispense the hot mixture (2% agarose-gelatine each, at 60°C) into the ice-cold mineral oil container (Supplementary Video S2). The drops of agarose-gelatine assumed a spherical shape and gradually sank to the bottom of the mineral oil while solidifying due to the low temperature. This manual method is effective if a small number of beads are required. We also developed a high-throughput system to improve efficiency and streamline the process. This involved using a magnetic stirrer with heating to maintain the solution in a homogeneous liquid state and employing a peristaltic pump to achieve a continuous flow of drops, enabling us to create thousands of beads with consistent sizes in a short period. We estimate that the above system can generate between 60 to 75 beads per minute. By standardising the drop volume to approximately 40 µl by modulating the pump flow rate, we achieved an average diameter of approximately 4 mm, ensuring a highly reproducible procedure (see Fig. 1, Fig. S1, Supplementary Video S1 and Table S1 for further details). The coefficient of variation remained below 2% across five independent batches (Figure S3, Table S1). This approach enabled the bulk creation of agarose-gelatine beads (AGBs), a typical size of glass beads used for biofilm cultivation [34]. The crosslink of the beads using DVS is a method akin to fabricating Sepharose, a chromatographic matrix commonly used for biomolecule purification [20]. Our protocol also demonstrated high reproducibility, as shown in Fig. S1, and the biogel showed significant stiffness and thermostability properties. Unlike its non-crosslinked counterpart, this biogel can withstand autoclaving without structural degradation. We also found that the beads need to remain in liquid throughout the production process and later for long-term storage since desiccation and later reconstitution are disruptive for the structure in contrast to chromatographic media. When possible, we tried to keep beads at the relevant medium for a specific experiment, starting from the autoclave step in that medium.

Next, we characterised the stability of the beads, which remained stable for at least five months after crosslinking and autoclaving. During the storage period in the culture medium, the AGBs did not change the aspect or diameter (Fig. 1F-G). The crosslinking not only provided the chemical stability to allow the beads to be autoclaved, but we also determined that the crosslinking significantly increased the beads’ strength, manifested by a significant increase in stiffness as measured by Young’s modulus of an atomic force microscope (Fig. 1H).

### *E. coli, P. aeruginosa* and *S. aureus* form robust biofilms on the agarose-gelatine beads

*E. coli*, *P. aeruginosa,* and *S. aureus* all formed biofilms on the surface of the AGBs, which were compared to glass beads. Quantification of bacterial biofilm cells by counting colony-forming units per millilitre (CFU/ml) revealed significantly more biomass for bacterial cells grown on AGBs than on glass beads for all strains (Fig. 2). We measured these values during the maturation process of biofilm at 24 and 48 hours. The number of CFUs for all three bacterial species was significantly higher for AGBs compared to glass beads at 24 and 48 hours. In preliminary experiments, we observed a higher variation of the bacterial count coming from the AGBs than from the glass beads. We suspected the heterogeneity was due to protocol limitations, so we modified the bacterial recovery protocol by incorporating mechanical and chemical methods. The method included using some enzymes to digest the extracellular matrix (see M&M and SM). We fundamentally used DNase I and papain, which helped in better dispersion and a good level of single cells in suspension without affecting bacterial viability, which is crucial for a satisfactory enumeration.

**Fig. 2.**
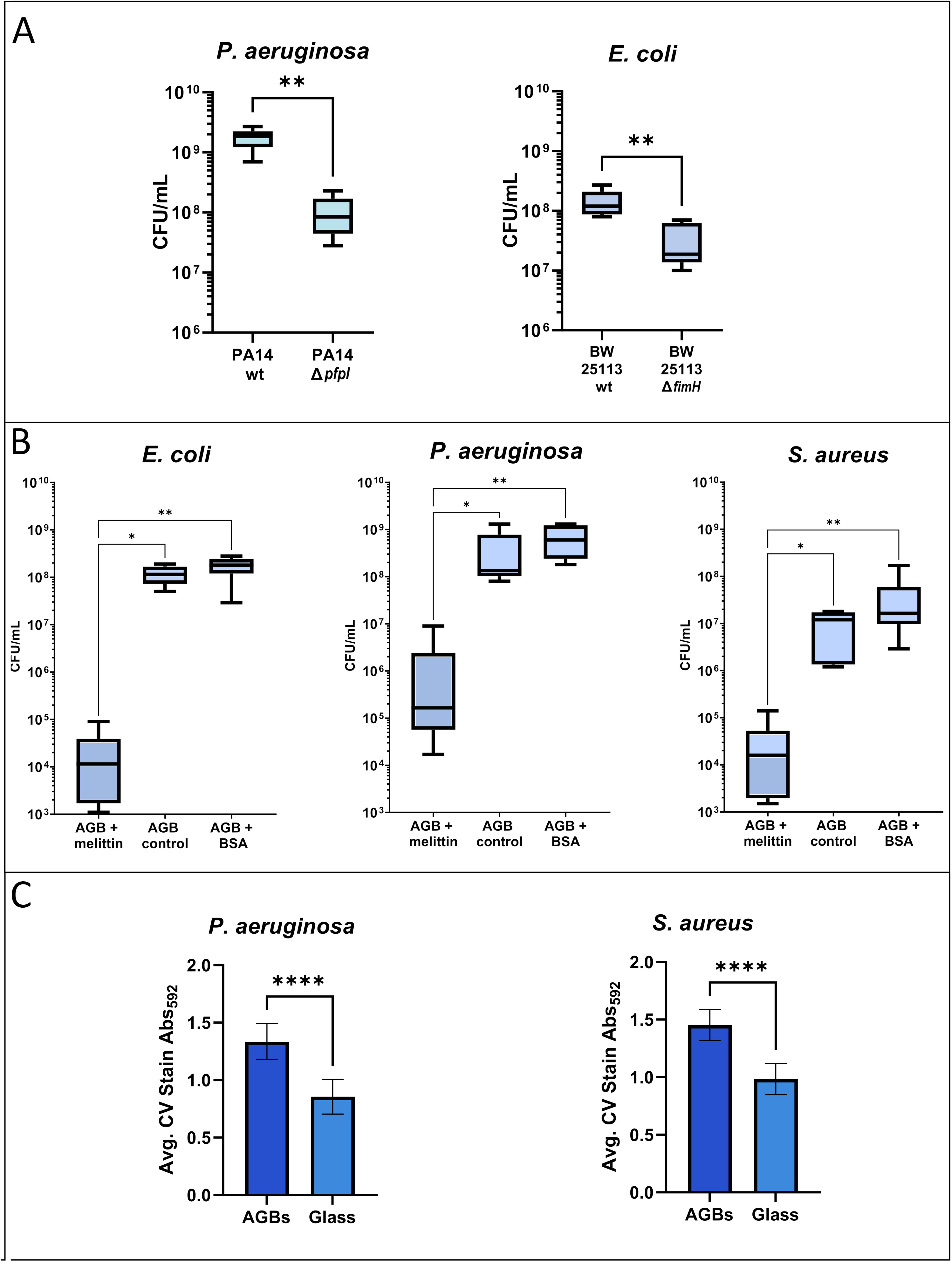
Comparative biofilm formation of E. coli, P. aeruginosa, and S. aureus in agarose-gelatine beads and glass after 24 and 48 hours. Boxplots of bacterial count from six biofilm-containing beads for each species grown on the surface of the beads at 24 and 48 hours (N=6). The samples were compared via the Wilcoxon-Mann-Whitney test, and the two asterix indicate that the p-values were below p<0.01. The Boxes extend from the 25 to the 75 percentiles, and the whiskers represent minimal and maximal values, the middle line representing the mean.

### Agarose-gelatine beads are compatible with scanning electron microscopy and live fluorescence microscopy

The transparency of agarose is particularly advantageous for live fluorescence microscopy, enabling clear visualisation of bacterial cells in real-time without interference. Furthermore, the biocompatibility of agarose-gelatine ensures that the bacterial cells remain viable and maintain their physiological states, leading to more reliable and relevant results in biofilm research. As a proof of principle, we constructed fluorescent strains of *E. coli*, *P. aeruginosa* and *S. aureus* on the surface of the AGBs using fluorescence confocal microscopy (Fig. 3A). We designed a 3D adapter (M&M, Fig. S4) for a more simplified and stable microscopy observation to study biofilms on 4mm AGBs (Fig. 3B).

**Fig. 3.**
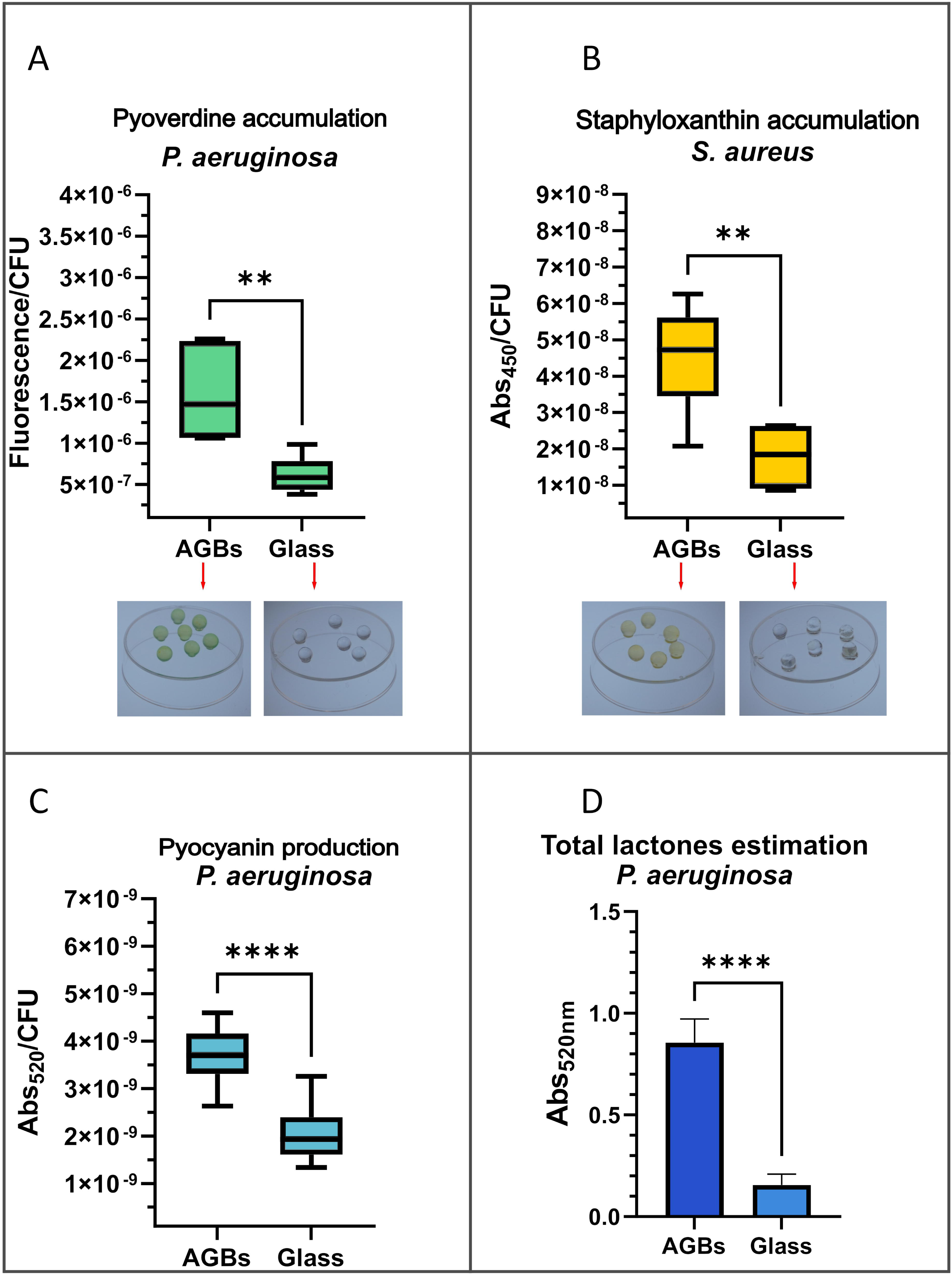
Microscopy of AGB biofilm formed by GFP-labelled E. coli, P. aeruginosa and S. aureus using fluorescence, confocal laser scanning microscopy and Scanning electron microscopy. AGBs were allowed to mature into a biofilm over 48 hours (A). A Z-stack comprising 29 images revealed a surface section of the AGB densely covered with an intact E. coli biofilm, and a single cross-section image extracted from the Z-stack displayed an AGB uniformly coated with E. coli biofilm. Later, the imaging process was made less tedious using a 3D printed adapter (See Material and Methods and Fig. S4), which also allows imaging with an oil immersion lens (B). After 48 hours of cultivation for E. coli, S. aureus and P. aeruginosa at different magnifications using Scanning Electron Microscopy, biofilms on the surface of an AGB (C).

Another relevant technique to study bacterial biofilms is scanning electron microscopy (SEM). Due to their hydrogel nature and spherical shape, the AGBs required special treatment (see M&M section) to fix, dehydrate, and keep the integrity of the structure of the beads and preserve the bacterial biofilm. The best results were obtained by drying all the beads with the critical point dryer. Then, we could perform scanning electron microscopy of *P. aeruginosa* and *S. aureus* biofilms (Fig. 3C). The gelatine component offers a suitable texture that facilitates sample preparation and enhances electron imaging, capturing fine details of biofilm architecture.

### Quantitative proteomics (liquid chromatography mass spectrometry, LC-MS) shows that biofilms on Agarose-Gelatine beads and glass beads have different physiology

Our investigation into the differences between bacterial biofilms growing on AGBs and glass beads, in terms of gene expression and bacterial physiological changes, was conducted with the rigorous method of label-free quantitative proteomics. This approach allowed us to identify distinct protein expression profiles in biofilms grown on agarose-gelatine beads (AGBs) compared to glass beads for three bacterial species - *E. coli*, *P. aeruginosa*, and *S. aureus*. It’s worth noting that we employed a modified protocol ad doc for this study with variations from previous works [35,36]. We identified and quantified 1113 proteins for *E. coli*, 1500 for *P. aeruginosa,* and 1251 for *S. aureus* (Supplementary Table S1, S2, and S3, respectively). Proteins showing at least a 2-fold change in their relative intensity between glass beads vs. agarose-gelatine beads, with an FDR-adjusted *p*-value < 0.05, were considered significantly changed. Significantly, we observed notable abundance changes in important virulence factors for *P. aeruginosa* and *S. aureus*, potentially indicating differences in surface sensing due to chemical composition (Fig. 4).

**Fig. 4.**
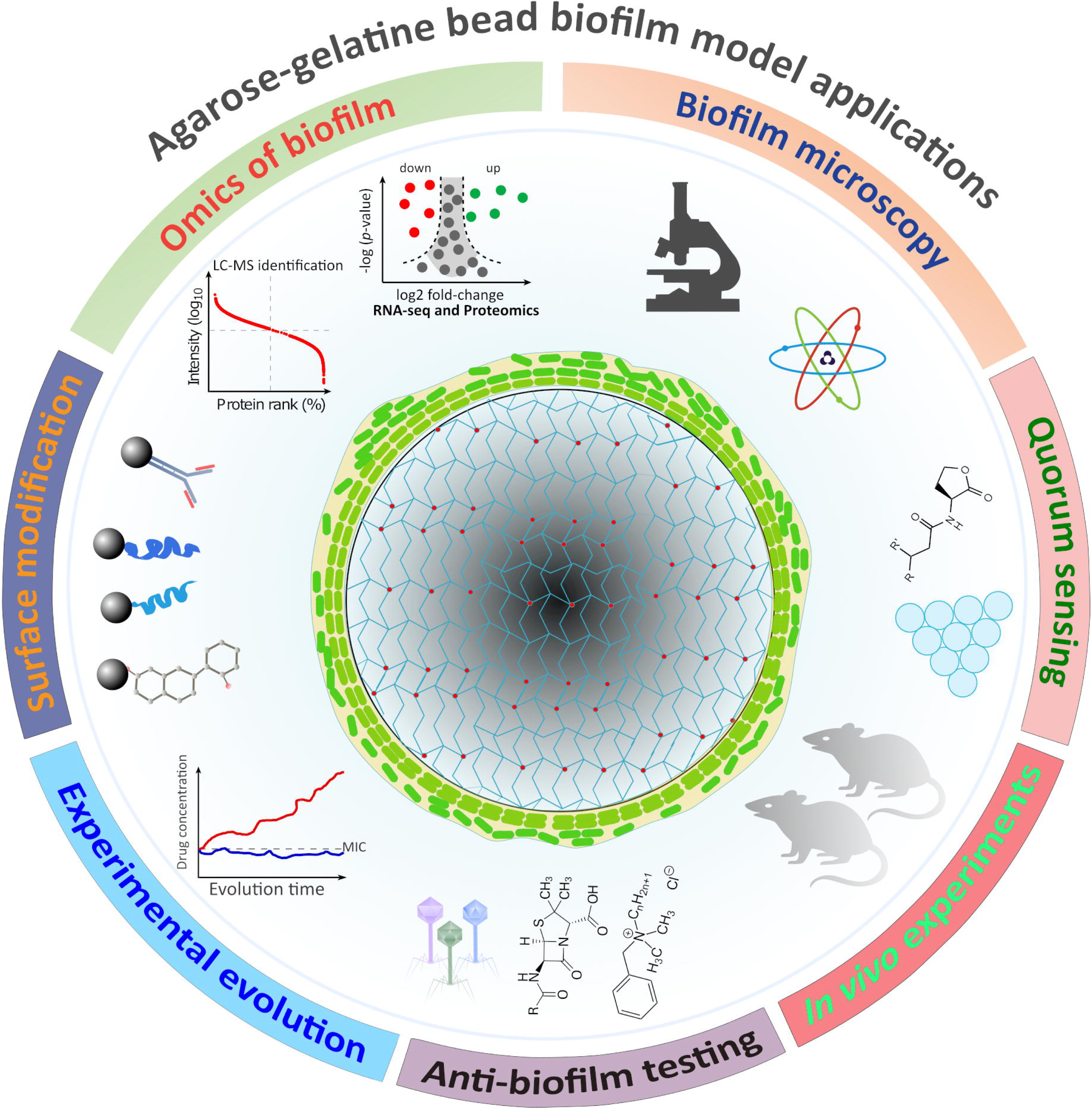
Comparison of relative wide proteome expression between agarose-gelatine beads and glass beads for bacterial biofilms of E. coli, P. aeruginosa, and S. aureus. Volcano plots show relative protein intensity changes between AGBs and glass beads measured by LC-MS and label-free quantification. Orange dots denote proteins significantly more abundant in AGB biofilms, while blue dots indicate proteins more abundant in glass beads biofilms. Proteins are considered significantly changed if they show at least a 2-fold change in their relative abundance (log2 fold change >1 or <-1) with an FDR adjusted p-value < 0.05, as indicated by the orange lines.

In the biofilms of *E. coli*, the 50S ribosomal subunit protein L35 (RpmI) had a noticeable increase in abundance when grown on AGBs compared to glass beads (log2 fold change (log2FC) = 2.5). Similarly, the small 30S ribosomal subunit S16 (RpsP) and ribosomal silencing factor (RsfS) were upregulated, but to a lesser extent. RsfS is known to be expressed in slowly growing cells and inhibits translation, indicating that the population of AGBs is in the mature phase [37]. On glass beads, there is a distinct upregulation of 50S ribosomal subunits RpmH and RplU, or subunits L34 and L21; at log2FC = −5.2, RpmH was the most differentially abundant protein. Apart from this striking difference in ribosomal structural content, several other ribosome-associated proteins were upregulated on glass: RsmH, Sra, YgaM and Rmf. RsmH is 16S-rRNA-m4C1402-methyltransferase, ubiquitous in bacteria and eukaryotic organelles; it facilitates a modification of the ribosomal P translation initiation site, allowing different codon usage for more efficient ribosome biogenesis [38]. The physiological roles of Sra and YgaM are less known but considered regulatory [37]. The former tends to bind active 70S Ribosomes during stationary phase and maturation, and the latter provides ribosomes with an anchoring point to the membrane. Rmf, or Ribosome modulation factor, is essential for the reversible formation of 70S-90S-100S Ribosome Dimers in gammaproteobacteria [37]. There is a higher expression of NusA (log2FC = 2.9) on glass beads, which acts as a transcription terminator factor and chaperone of rRNA, increasing their numbers; NusA is expressed following stress such as cold shock [39]. Overall, we see different ways of maturing and diversifying the ribosomes in size, structure, activity, and biogenesis. This trend is evident in all three strains, with potentially more regulatory shifts and stress response adaptations in biofilms on glass beads. The data also shows a notable change in cell division via Z-Ring formation through FtsZ polymerisation, an essential process for cell elongation and division. Notably, ZipA, which anchors FtsZ to the membrane, was significantly more abundant (log2FC = 2.2) on AGBs, contrasting with the upregulation of ZapA (log2FC = −2.1) on glass beads, influencing cell division dynamics [40].

In *P. aeruginosa*, important virulence factors were found to be more abundant in AGBs than in glass beads. Elastase (LasB) showed the most significant change in relative abundance (log2FC = 5.1). Elastase is regulated by the LasR/LasI quorum-sensing system, indicating increased virulence factor activity in biofilms grown on AGBs [41,42]. PrpL (log2FC = 3.2), also known as Protease IV or lysyl-endopeptidase [41,42], which can cleave fibrinogen and disrupt the activity of the immune system, was also elevated. Several proteins involved in pyoverdine biosynthesis, including PvdA, PvdJ, PvdD, and PvdL, also showed an increased abundance in AGB-biofilms. Pyoverdine’s primary function is iron chelation from host cells, which is involved in *in vivo* infections [43]. Another significantly upregulated protein in AGBs is the aminopeptidase Lap, also known as PaAP, which cleaves extracellular proteins [42] and PhzM, involved in pyocyanin biosynthesis [44]. The only virulence-related upregulated protein on glass beads was AmrZ, also known as AlgZ (log2FC = 3.0). This important transcription factor regulates Alginate production and motility [45,46]. Conversely, among the downregulated proteins on AGBs were components of the Sec translocon system, such as YajC (log2FC = −4.4) and SecF (log2FC = −2.5), indicating reduced protein export and altered membrane dynamics [47]. The second and third most abundant proteins on glass beads are uncharacterised and coded by the genes PA0049 and PA5305 (with log2FC values of −4.2 and −3.7, respectively). PA0049 expression is under the stringent redundant response regulator DksA in *P. aeruginosa* [48]. A large proportion of proteins are probable, putative, uncharacterised, or hypothetical for *P. aeruginosa* (18 of the 55 abundant proteins on AGBs and 113 of the 240 on glass beads, between 30-50%), detected to be above our specified threshold for significant change. *S. aureus* showed a similar trend, making comparative analysis more intricate for both organisms relative to *E. coli*.

We observed a similar trend in *S. aureus* biofilms as in *P. aeruginosa*, where we found higher virulence factor expression in AGBs than in glass beads. However, *S. aureus* showed a more complex response with more differentially expressed proteins for both substrates (247 proteins on AGBs versus 297 on glass beads). *S. aureus* had about twice the number of proteins with altered levels compared to *E. coli* and *P. aeruginosa*. These results indicate that different species have differential responses in the class of proteins and the number of genes that respond to the surface where the biofilms develop.

DeoC was the highest expressed protein on AGBs (log2FC = 5.0), and it is usually part of the inducible *deo* operon in bacteria, enabling the metabolism of exogenous nucleosides [49]. DeoC, or deoxyribose phosphate aldolase, is a duplicated gene in *S. aureus*, and both copies are highly upregulated on AGBs (log2FC = 2.5), along with other genes in this operon, DeoB and DeoD [50]. The second most abundant protein on AGBs was MsrB (4.9 log2FC), a methionine sulfoxide reductase that protects against oxidative stress and is a marker for mature biofilms [51]. DltC, critical in teichoic acid biosynthesis, cell envelope overall charge, and virulence, was next on the list [52]. The most abundant protein on glass beads was SAOUHSC_02161 (8.0 log2FC), an MHC class II analogue protein of unknown function, under the SaeRS two-component system regulon and considered an attachment protein [53].

Several transcription factors and additional regulators were significantly increased on glass: SarR, FapR, Rex, RecX, MsrR, and Spx; while on AGBs, seven other transcription factors were upregulated: ArcR, NusB, NusG, CodY, NusA, SrrA, RsbW and VraR. SarR downregulates the expression of SarA, which usually activates the *agr* operon quorum sensing system, leading to increased production of exotoxins required for virulence [54]. FapR is a regulator of membrane lipid biosynthesis and lipid homeostasis [55]. Rex is a redox-responsive repressor activated by the ratios of NADH:NAD in cells; its regulon enzymes are involved in many metabolic pathways [56]. RecX is an anti-recombinase as it inhibits RecA under accumulated DNA damage [57]. MsrR regulates the cell envelope homeostasis and virulence factor production by structural means, abundant in antibiotic-tolerant cells [58]. Spx is a global effector that accumulates cells during various stresses and regulates many pathways, most notably thioredoxin, by a unique mechanism [59]. In biofilms on AGBs, the most abundant regulator is ArcR, which is critical for anaerobic metabolism, mainly allowing the metabolism of arginine over sugars [60]. Nus is an accessory protein forming a complex with RNA polymerase, enhancing bacterial rRNA transcription and synthesis [61]. CodY regulates the *agr* virulence system depending on population density and nutrient scarcity [62]. SrrA is part of the two-component system SrrAB that increases virulence factor production through *agr* at low oxygen levels [54]; RsbW is a serine-protein kinase acting as an anti-sigma factor to SigB, inhibiting transcription after binding [63]; and VraR is part of the two-component system VraSR, which is upregulated in response to cell-wall damage affecting peptidoglycan homeostasis [64].

Regarding other pathways leading to virulence factor abundance, we observed a differential upregulation on glass beads, notably the biosynthesis pathway of staphyloxanthin: AldH1, CrtM, CrtP, and CrtN. Additional virulence factors that were more abundant on glass beads comprise two fibrinogen-binding proteins (SAOUHSC_01114, SAOUHSC_01110), immunoglobulin binding Sbi and two uncharacterised leukocidin-like proteins (SAOUHSC_02241, SAOUHSC_02243). Several subunits of Phenol-Soluble Modulin are upregulated (psmA1/A2 and psmA4). ClfA, or clumping factor, facilitates cells’ clumping, thus defending them from phagocytosis. There is a notable abundance of extracellular active enzymes such as serine protease SplE and Coagulase (SAOUHSC_00192) and a plethora of known staphylococcal antigens (IsaB, SsA2, SsL10, SsaA). On AGBs, we can detect the upregulation of virulence-promoting factors, such as TraP and LuxS. TraP is involved in the upregulation of the *agr* operon in a cell density-dependent manner [65], LuxS is the enzyme which produces autoinducer-2 molecule, which in turn results in enhanced biofilm phenotypes [66]. Other notable enzymes involved in pathogenesis were also more abundant on AGBs, including lipase-1 LipA, catalase katA, and components of the type VII secretion system components, EsxA and EsaG, important for intracellular infection of the host [67].

### The AGBs are suitable for the detection of biofilm-deficient strains

Our model system presents a valuable tool for detecting or screening genes and conditions influencing biofilm formation. To validate this concept and assess the capability of AGBs to detect variations in biofilm formation, we examined two bacterial strains known for their deficiencies in biofilm formation. Previous biofilm research has shown that type I fimbriae, produced by the *fim* operon, are crucial for forming submerged biofilms. Additionally, type I pili, including the mannose-specific adhesin FimH, are for initial surface attachment [68]. Earlier studies reported that disrupting the *pfpI* gene in *P. aeruginosa* decreased the bacterium’s ability to form biofilms [69]. When we cultivated both mutant strains in AGBs and compared them with their respective parental wild-type strains, we observed a significant decrease in biofilm formation (Fig. 5A). This finding indicates that our AGB system is suitable for detecting differences in biofilm formation between strains.

**Fig. 5.**
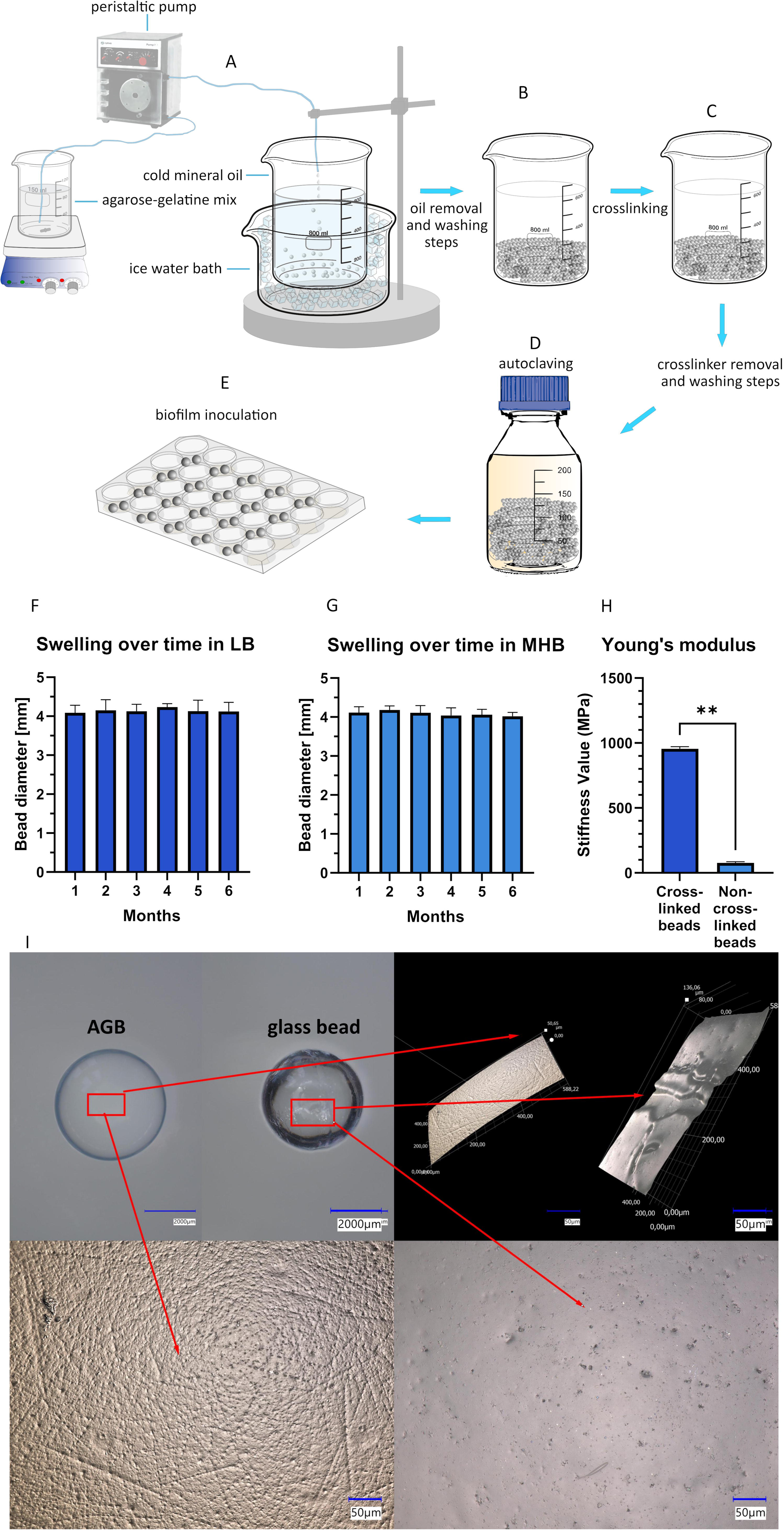
Diverse experiments with biofilm formation, including biofilm deficient strains, antibiofilm surface treatment and crystal violet staining. (A) Two biofilm-deficient strains of E. coli (ΔfimH) and P. aeruginosa (ΔpfpI) on AGBs after 24 and 48 hours. Boxplots of bacterial count from biofilms of E. coli and P. aeruginosa mutants compared to their respective parental wild-type strains (N=6). (B) Comparative biofilm formation of E. coli, P. aeruginosa and S. aureus on melittin-decorated agarose-gelatine beads after 48 hours of incubation. Boxplots of bacterial counts from biofilms attached to melittin-coupled, uncoupled, and BSA-coupled after 48 hours of incubation (N=6). (C) Biofilm formation of P. aeruginosa and S. aureus on AGBs vs. glass beads after 48 hours of incubation using the modified crystal violet staining protocol. Bar charts represent the average absorption of the dye at 592nm that remained bound to the biofilms, a qualitative measure of biofilm formation capacity. We measured ten repetitions for each species on each substratum (N=10). The samples were compared via the Wilcoxon-Mann-Whitney test. One asterisk represents p<0.05, two asterisks for p<0.01, three for p<0.001 and four for p<0.0001.

### Chemical modification of agarose-gelatine beads is a valuable strategy for investigating anti-biofilm compounds

One of the significant advantages of crosslinked agarose is its potential for conjugating diverse chemical compounds to its surface. This feature benefits our biofilm model and has practical implications for investigating antibiofilm compounds to protect surfaces, a crucial aspect of biofilm research. To demonstrate this concept, we chose melittin, an antimicrobial peptide known for its anti-biofilm properties [70]. Following the protocol described by March *et al*., we used the cyanogen bromide method to couple melittin to agarose-gelatine beads [71]. Cultivating biofilms of our three bacterial models on melittin-decorated AGBs revealed a dramatic decrease in bacterial attachment to the bead surfaces compared to untreated and BSA-coupled beads. For all three bacterial species—*E. coli*, *P. aeruginosa*, and *S. aureus*—the number of biofilm-forming bacteria was reduced by 4 to 5 orders of magnitude compared to the controls (Fig. 5B). Biofilms grown on BSA-coupled beads exhibited the highest biofilm formation, indicating that the chemicals used for protein coupling to the AGBs were not responsible for the observed biofilm inhibition. Notably, final counts of planktonic bacteria did not differ in wells where bacterial species were incubated, indicating that the observed effects were specific to biofilm formation.

### Agarose-gelatine bead biofilms can be quantified with a crystal violet assay

A popular and convenient method for quantifying bacterial biofilms is to measure the amount of crystal violet stain retained by the biofilms [72]. We tested this method with our AGBs and compared the performance of the crystal violet staining with biofilms grown on glass beads. Initially, when using the standard protocol, we observed that the beads accumulated a significant amount of background crystal violet compared to the glass beads. To address this issue, we made some modifications to the protocol. By adjusting the crystal violet concentration, incubation temperature, and washing times, we optimised the protocol for the AGBs. As a result, we confirmed through the crystal violet assay (see Methods and Materials section) that bacteria growing on the surface of AGBs form more biofilm than their glass bead counterparts (Fig. 5C).

### Agarose-gelatine beads can serve as small molecule reservoirs

In contrast to other abiotic biofilm substrates, the permeability of crosslinked agarose could serve as a reservoir of small molecules such as nutrients, quorum-sensing molecules, drugs, etc. This property can be advantageous in certain experiments as it may occur in *in vivo* biofilms where bacteria are attached to host surfaces, such as mucosal layers or necrotic tissue debris. Our experiments involve two bacterial models that produce detectable substances that could accumulate within the agarose beads. We observed that the agarose beads consistently appeared yellower for *S. aureus* or greener for *P. aeruginosa*, indicating the diffusion of pigments into the inner part of the beads (Fig. 6A-B). For *P. aeruginosa*, upregulation of the biosynthetic pathway of Pyoverdine was evident in AGBs; in contrast, upregulation of the Staphyloxanthin was apparent on glass beads. To try to quantify changes in the concentration of those pigments in bead-containing samples, we adapted some chemical and physical protocols (Materials and Methods) to extract them into the solution that could be measured with a plate reader relative to CFU/mL numbers (Fig. 6A-C); A similar approach to this protocol has been taken in a recent publication [73]. We also propose that this hydrogel could facilitate the diffusion of soluble molecules, including nutrients and bacterial quorum-sensing molecules while retaining some molecules within its structure as a reservoir, as observed in many *in vivo* systems. We estimated the quantity of N-acyl-homoserine-lactones (AHL) in the *P. aeruginosa* PAO1 bead-containing samples using a previously described colourimetric method [74] with adaptations. The assay indicates a significantly higher lactone content in AGB samples than glass (Fig. 6D).

**Fig. 6.**
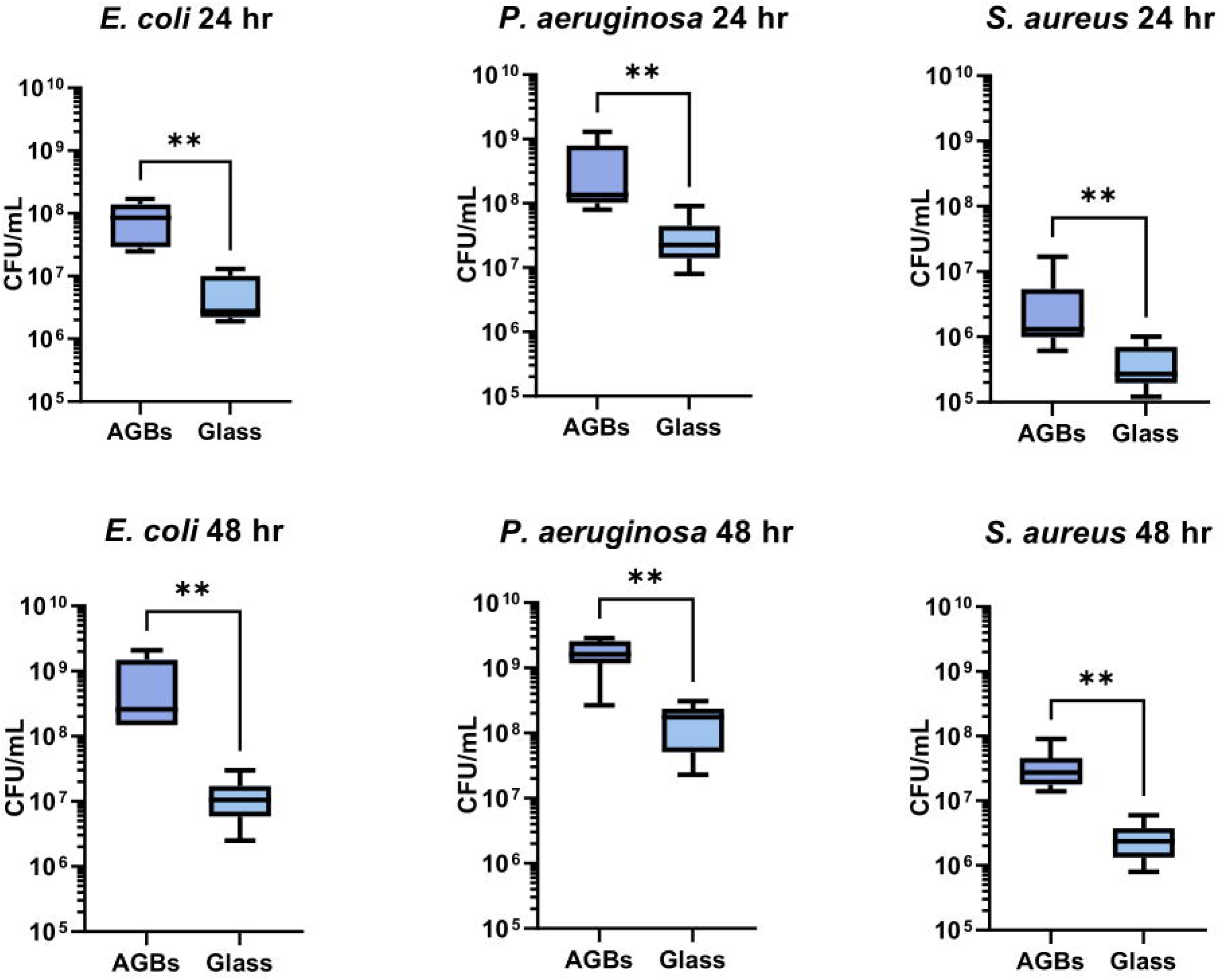
Comparative amounts of different virulence factors from P. aeruginosa and S. aureus on agarose-gelatine beads after 48 hours of incubation. Boxplots of (A) pyoverdine, (C) pyocyanin (produced by P. aeruginosa), and (B) staphyloxanthin (produced by S. aureus) accumulation from biofilms after 48 hours of incubation (N=6). (D) Estimating N-Acyl homoserine lactones in P. aeruginosa shows the possible effect of diffused molecules on AGBs on quorum sensing regulated activity. The samples were compared via the Wilcoxon-Mann-Whitney test. One asterisk represents p<0.05, two asterisks for p<0.01, three for p<0.001 and four for p<0.0001. The Boxes extend from the 25 to the 75 percentiles, and the whiskers represent minimal and maximal values, with the middle line representing the mean.

## Discussion

Our biofilm model has significant potential for a wide range of research applications (see Fig. 7). AGBs can be used to test anti-biofilm molecules, evaluate surface protection compounds, study quorum sensing mechanisms, investigate biofilms on foreign bodies, and conduct microscopy and ‘omics profiling studies of biofilms. Additionally, our system is compatible with traditional experimental approaches, including crystal violet quantification and live and scanning electron microscopy.

**Fig. 7.**
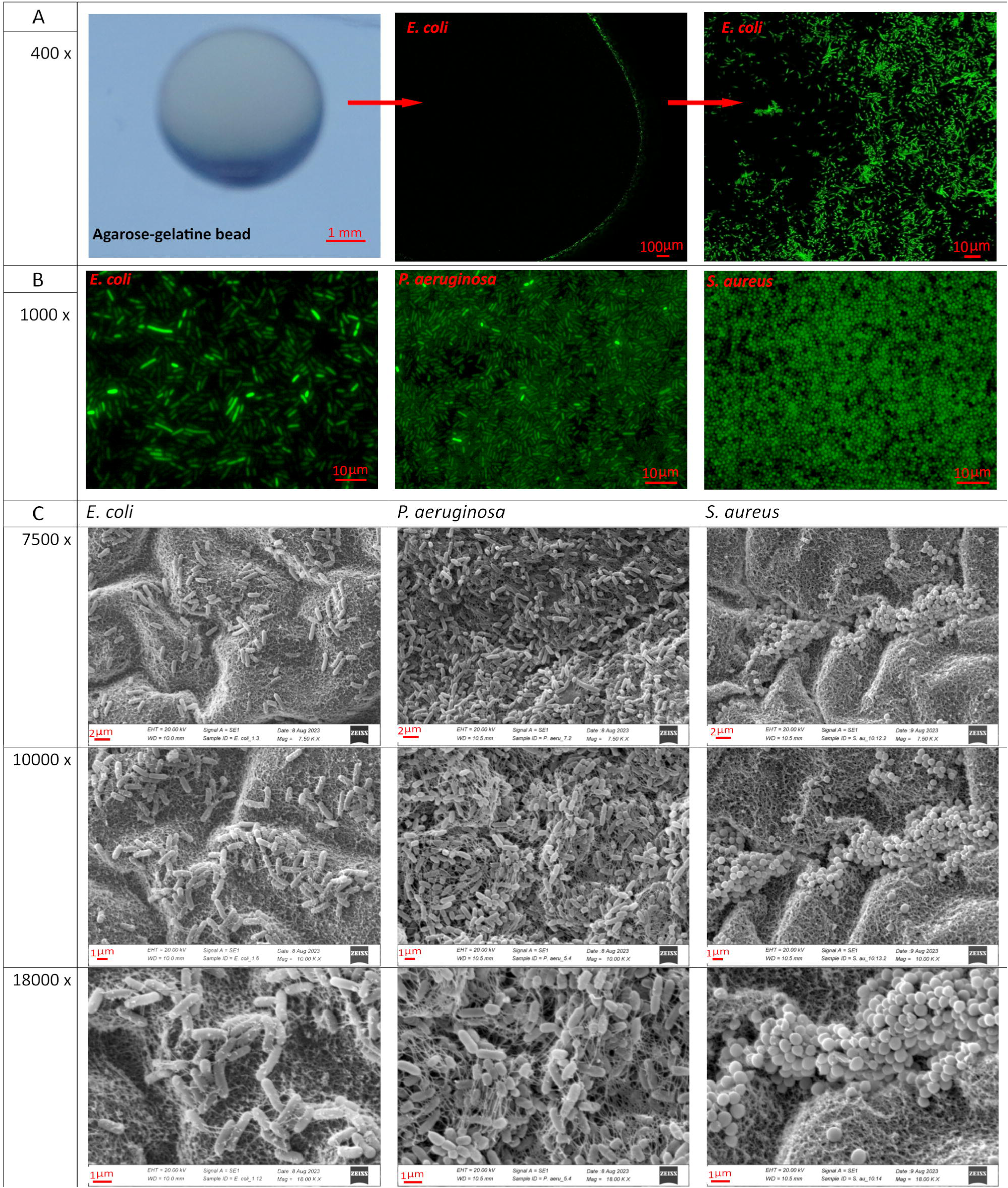
Summary of the potential applications of the new model of crosslinked agarose-gelatine beads. These beads, with their biocompatibility, ease of preparation, and versatile physical properties, hold significant potential as substrates for biofilm-related research and applications. Their adaptability to mimic various environmental conditions, from nutrient-rich to nutrient-limited, provides a versatile platform for studying biofilms under different scenarios. They can be used to investigate antibiofilm agents, evaluate the efficacy of various compounds in disrupting or inhibiting biofilm formation, and develop new antibacterial treatments and coatings. Moreover, these beads can facilitate studying quorum sensing (QS) mechanisms within biofilms. Furthermore, the beads are compatible with various analytical techniques, including live fluorescence and scanning electron microscopy.

One attractive application of our system is the ease of conjugating molecules to the surface of the beads. The chemical modification of the AGBs is a great advantage, as demonstrated in this study by conjugation with melittin. The attachment of this antimicrobial and antibiofilm peptide significantly reduced biofilm formation by *E. coli*, *P. aeruginosa*, and *S. aureus*. Melittin, a small cationic peptide composed of 26 amino acid residues [75], is a principal constituent of bee venom and is used as a tool of social immunity to protect their colony from pathogen invasion [76]. This property of easy compound conjugation can also be used to investigate what elements stimulate biofilm attachment and avoid certain materials or use them in beneficial biofilms [77].

Agarose-gelatine beads exhibit a unique capability to serve as reservoirs for small molecules. They allow for the gradual release of nutrients, quorum-sensing molecules, drugs, and other minor compounds, thus providing a more accurate simulation of *in vivo* biofilm conditions. This feature is advantageous for experimental setups replicating complex environments where bacteria adhere to host surfaces or necrotic tissue debris [78]. The diffusion of virulence factors such as staphyloxanthin, pyoverdine, pyocyanin, and quorum-sensing molecules exemplified this capability. Staphyloxanthin is an interesting example because the proteomic analysis shows downregulation of its biosynthesis. It is present in cell membranes, protecting lipids, proteins, and DNA from ROS [79]. The main enzymes in ROS defence are catalases and superoxide dismutase, while staphyloxanthin provides additional protection [80]. Clinical samples from diabetic patients’ foot ulcers have shown that staphyloxanthin impairs wound healing and enhances *S. aureus* survival in presence of ROS [81]. It is known that a substantial reduction in staphyloxanthin production occurs in correlation with biofilm maturation [82]. Additionally, staphyloxanthin accumulation within the AGBs, as we have shown, could be responsible for inhibiting pigment production.

The biofilm system based on AGBs proves effective for detecting genetic factors influencing biofilm formation, as demonstrated by studying mutant strains deficient in biofilm formation, explicitly focusing on the *fimH* of *E. coli* [83] and the *pfpI* gene in *P. aeruginosa* [69]. Comparing biofilm formation in AGBs between mutant and wild-type strains revealed a significant reduction in the mutants’ biofilm formation, validating the AGB system’s capability to detect variations in biofilm formation under controlled conditions.

The quantitative proteomic profiling performed in this study revealed distinct protein expression profiles for *E. coli*, *P. aeruginosa*, and *S. aureus* biofilms grown on AGBs compared to glass beads. Some studies suggest that bacterial growth on abiotic surfaces significantly differs from the bacteria from *in vivo* biofilm-related infections [84]. Many more differentially expressed proteins are crucial virulence factors for *P. aeruginosa* and *S. aureus*, likely reflecting differences in surface sensing. For *E. coli*, there were significant differences in ribosomal protein expression in biofilms on AGBs and glass; A similar trend was observed for *P. aeruginosa* and *S. aureus*. In *P. aeruginosa*, essential virulence factors such as elastase (LasB) and the pyoverdine synthesis pathway were more abundant in biofilms on AGBs. For *S. aureus*, biofilms on AGBs and glass beads exhibited differences in protein expression and gene regulation; it exhibits differential expression of factors for host cell recognition and utilisation of host resources. The study identified several regulatory proteins that mediate the expression and effects of the quorum sensing system. In conclusion, biofilms from AGBs and glass beads represent different phenotypes, with the former possibly resembling a mature and established biofilm and the latter resembling a phenotype closer to a bloodstream infection.

It does not escape our attention that this new biofilm model offers several applications. Firstly, it may enable *in vivo* experiments that investigate biofilms in the context of infections associated with foreign objects, providing crucial insights into the dynamics and treatment of such infections. Secondly, the hydrogel material could be potentially used in bioengineering and 3D printing as a biopolymer for fabrication, opening avenues for advanced tissue engineering and regenerative medicine. Lastly, AGBs can serve as an effective model for understanding the role of biofilms in bacterial resistance to antibiotics, allowing the scientific community to conduct experimental evolution studies that could lead to novel strategies for combating antibiotic resistance and biofilm-associated infections. Besides, because of the hydrogel nature of the AGBs, there is also a potential utilisation to study liquid air interface biofilm.

## Conclusions

In this study, we have introduced a new substrate based on crosslinked agarose-gelatine hydrogel in the form of beads as a novel strategy to investigate biofilms of bacterial pathogens. We have shown that this method is compatible with various biofilm structural studies and quantitative techniques, including live microscopy techniques such as fluorescence and confocal microscopy, as well as scanning electron microscopy. Our model is also compatible with other biofilm quantitative techniques, including crystal violet staining, and we have developed a protocol for the enumeration of viable cells, which maximises recovery and dispersion using enzymatic and mechanical disruption of the biofilm. The agarose-gelatine bead model provides a new opportunity to study biofilm physiology, as the permeability of the hydrogel beads allows the diffusion of small molecules, including drugs, nutrients, quorum sensing signalling molecules, and other substances such as siderophores and other secretion products. We have demonstrated that the beads maintain their properties for an extended period, at least several months.

## Materials and Methods

### Generation of agarose-gelatine beads

To produce the bead substrate, 4 g of each agarose and gelatine were added to 200 ml (2% each w/v) of distilled water in a 400 ml bottle and autoclaved for 20 minutes at 121°C and 1 atmosphere. After sterilisation and throughout the production process, the liquid mixture was kept liquid by placing the flask on a hot plate of the stirrer at 65°C with moderate mixing. In a separate container, 500 ml of mineral oil (Merck, Germany) was placed in an ice water bath. The oil was previously degasified to remove air bubbles using a vacuum pump. We used two procedures to produce the agarose-gelatine beads. First, we used a multistep pipette (Eppendorf, Germany) to spot around 40 µl of substrate in the cold mineral oil, ensuring that the tip was between 1 to 2 cm from the surface of the oil to avoid deformation of the beads while solidifying or partitioning of the drops (See Fig. S1). A second and higher throughput method incorporated a peristaltic pump P-1 (Pharmacia Biotech, Sweden) that allows a continuous and semiautomatic dropping of the beads into the mineral oil (Fig. 1). At the end of the silicon tubing, a 2 mm internal diameter stainless steel capillary was incorporated for a more precise holding and stable drops, two silicone tubes with a diameter of 3 mm and a length of 0.5 m were extended with one metal tube of the same diameter and a length of 20 cm. The flexible ends were connected to the peristaltic pump, while the ends with the metal tubes opened into the bottle with the substrate or above the collection beaker, respectively. The movement of the peristaltic pump resulted in a stable output of uniform drops of the agarose-gelatine mix, which solidified upon entering the inert, cooled mineral oil, forming ~4 mm Agarose-gelatine beads (AGBs). After processing, the beads were sieved and washed with distilled water with at least 50 volumes until the oil was removed, allowing them to rest for five minutes between washes to allow the oil separation from the aqueous phase.

### Crosslinking of the agarose-gelatine beads

The washed and clean agarose beads were crosslinked using divinyl sulfone (DVS) using a modified method from the original described elsewhere [21]. The beads were weighed to calculate the required amount of DVS (v/w) (Merck, Germany). Approximately 500 ml of 0.5 M sodium carbonate buffer, pH 11, was used per 100 grams of gel beads (five volumes). The beads were gently transferred with the help of a funnel to an 800 ml flask, capped, and the crosslinking reaction was initiated by adding DVS, a final concentration of 0.5%. The reaction was allowed to proceed on a shaker at room temperature overnight, covering the beaker with parafilm to prevent evaporation. The next day, the beads were thoroughly washed with distilled water by decanting and sieving until a neutral pH was achieved. To remove residual amounts of unreacted vinyl groups, we placed the beads in five volumes of 0.5 M sodium carbonate buffer, pH 9, and added beta-mercaptoethanol for a final concentration of 0.5%. The flask was incubated with shaking overnight in the dark at room temperature. Afterwards, 500 ml of distilled water was used to wash the beads five times. To remove the residual unreacted vinyl group, the beads were covered with five volumes of 1 M sodium carbonate, pH 10, and glycine was added for a final concentration of 10% (w/v). The reaction was also conducted overnight at room temperature with shaking. The next day, the beads were washed three times with five volumes of 0.1 M carbonate-bicarbonate buffer at pH 10 and a 0.1 M citrate buffer at pH 4.0. The beads were rewashed with five volumes of distilled water three times and added directly to a flask containing five volumes of bacterial culture media lysogeny broth (LB) or Mueller-Hinton broth (MHB). The bottles were autoclaved and stored at +4°C until use. For a step-by-step protocol, see Protocol I of supplementary materials.

### Microscopy of Bead Surfaces

To visualise the intact surfaces of agarose-gelatine glass and beads, we utilised a VHX-X1 microscope (Keyence, Austria). Glass beads were directly placed into a 45 mm petri dish for observation. AGBs were first removed from the preservation medium (LB). Then, the excess liquid was carefully pipetted off before placing the beads into a separate 45 mm petri dish for direct examination at different magnifications.

### Bacterial strains and cultivation conditions. Escherichia coli

K-12 MG1655 [85], *Escherichia coli* BW25113, and its derivative mutant *E. coli* BW25113 *fimH::*Kn, both from the Keio collection [86], *P. aeruginosa* PAO1 [87], *P. aeruginosa PA14* [88] *and its derivative mutant P. aeruginosa PA14 pfpI::*MAR2xT7[88], and *Staphylococcus aureus* SH1000 [89] were the strains used for different experiments across this study. Generally, bacteria were cultivated on Petri dishes from frozen glycerol stocks on lysogeny broth (LB) agar and incubated overnight at 37°C. The next day, a single colony was inoculated into 10 ml of liquid LB in 50 ml Falcon tubes and shaken at 200 r.p.m. overnight at 37°C. The total cell density was determined by measuring the optical density (OD) in a spectrophotometer at 600 nm. The inoculums were diluted to approximately 10⁶ CFU/ml per culture for biofilm inoculation.

### Biofilm cultivation

For biofilm culturing, two beads were aseptically transferred to each well of a non-binding polypropylene 24-well microplate (Carl Roth, Germany). Overnight cultures of *P. aeruginosa, S. aureus* SH1000, and *E. coli* K-12 MG1655 in LB medium were diluted to an optical density of approximately 1×10^5^ CFU/ml and added to the microplate containing the beads (1 ml per well). The 24-well microplate was placed in a multiwell plate shaker with moderate shaking (20-30 r.p.m.) at 37°C for 24 and 48 hours. After 24 hours of incubation, the beads were gently washed with sterile 0.9% NaCl saline solution by pipetting, and a fresh medium was added.

### Biofilm viable bacterial enumeration

After biofilm cultivation, each bead was transferred with sterilised tweezers into a 1.5 ml microcentrifuge tube (Eppendorf, Germany) containing 1 ml saline. The samples were sonicated in an ultrasonic bath to detach the biofilm from the bead surface and separate the bacteria at 40 kHz for 3 min. Before and after sonicating, each microcentrifuge tube contained a bead and a vortex vigorously mixed with the saline solution. The tubes were centrifuged at 10,000 x g for 3 minutes, and the pellets resuspended in 200 μl of digestion buffer (150 mM NaCl, 50 mM Tris-buffer, 10 mM MgCl2, 5 mM L-cysteine, 5 mg/ml DNAse I, 2 mg/ml RNAse A, pH 7.5) and incubated at 37°C for 15 minutes. Afterwards, 20 μl of papain stock solution (10 mg/ml) was added to each tube and incubated for another 15 minutes at 37°C in moderate shaking. The tubes were centrifuged at 10,000 x g for 3 minutes and resuspended in 1 ml of 0.9% NaCl. A volume of 20 μl of each sample was transferred to the wells of the first row of a 96-well microplate. Using a multichannel pipette, all samples were diluted in 1:10 steps (20 μl sample; 180 μl saline) from the first row to the last. Droplets of 10 μl for each dilution were spotted on pre-dried LB agar plates (for 45 minutes before use in a laminar flow cabinet). The wells containing the bacterial dilution were carefully mixed by pipetting before spotting the drops. After drying the droplets (approximately 5– 10 minutes), plates were incubated for 16 hours at 30°C. The next day, the microcolonies were observed and, if necessary, allowed to grow for another two hours at 37°C. Then, the colonies were counted for the first dilution, which resulted in a countable range of 5–35 colonies. The bacterial number was determined, considering the dilution factor, and expressed as CFU/ml.

### GFP-tagging

To perform live fluorescent microscopy of bacterial biofilms on the surface of agarose-gelatin beads, a constitutive green fluorescent protein (GFP) gene was inserted into the chromosomes of *E. coli* MG1655, *P. aeruginosa* PAO1 and *S. aureus* SH1000 using ADDGENE Plasmids (pGRG36-Kn-PA1-GFP, pMF230 and pTH100 accordingly) and following the relevant methodologies published elsewhere [90–92]. Briefly, *E. coli* DH5alpha were used as plasmid donors carrying either the pGRG36-Kn-PA1-GFP or the pMF230, and *E. coli* DC10b was used for plasmid pTH100 for integrating GFP expression cassette into the chromosome of the strains used in this research accordingly. Donor strains were grown under specified conditions using LB broth and relevant antibiotics. Plasmids were purified using the Pure Yield Plasmid Miniprep System (Promega, USA) following the manufacturer’s instructions and stored at −20°C until use. Mid-exponential cultures (100 ml) of *E. coli* MG1655, *P. aeruginosa* PAO1, and *S. aureus* SH1000 were grown until an OD600 between 0.4 and 0.6 was reached. *E. coli* and *S. aureus* acceptors were made electrocompetent using previously described protocols [93,94] transformed with the relevant plasmids using electroporation (Gene Pulser Xcell™, Biorad, USA) using the manufacturer’s preset protocols for *E. coli* and *S. aureus*. *P. aeruginosa* was transformed by triparental conjugation, essentially as described [91] using the helper strain *E. coli* S17-1*-lambda-pir* carrying pRK2013 [95]. Growth on LB Agar and a fluorescence-adapted stereomicroscope checked the phenotypes of marker antibiotic resistance and green fluorescence, respectively. Positive clones were selected, grown, and kept as glycerol stocks at −80°C until use.

### Crystal violet staining

Qualitative quantitation of all biofilm components was performed using crystal violet staining [96]. For the substrate comparison, AGBs (4 mm) and glass beads (4 mm, Merck, Germany) were inoculated with *P. aeruginosa* PAO1 and *S. aureus* SH1000. After the maturation of the biofilm in 24-well polypropylene plates (Carl Roth, Germany) and a final wash, the beads in each well were covered with 500 μl of NaCl 0.9% saline solution and another 500 μl of a 0.1% crystal violet (Merck, Germany) in saline for 1 min. Using a single-channel pipette, the staining solution was aspirated and discarded. The beads were carefully rinsed by adding 1 ml of 45°C prewarmed distilled water several times, with 5 min incubation time between washes. The washing process was repeated until the water was clear or the wash had an OD592 below 0.01. To solubilise crystal violet retained by the biofilm, 1 ml of a mixed solution of 95% Ethanol [96] and 0.05% of Triton X-100 [97] were added to each well. The beads were incubated with gentle shaking for 30 minutes at 45°C. 200 µL of each sample were then transferred to the wells of a 96-well plate. The amount of crystal violet was estimated by absorbance at 592 nm.

### Scanning electron microscopy

To visualise biofilm formation on the surface of the bead substrate, images of colonised Agarose-gelatine beads were taken using a scanning electron microscope (SEM) (EVO MA 10, Carl Zeiss, Germany). Before removal from the 24-well plates where the biofilm developed, the samples were washed with saline, as described above. The following fixation steps were carried out under the safety cabinet due to the high toxicity of the chemicals used. Six samples of mature biofilm produced by *E. coli* K12, *P. aeruginosa* PAO1 and *S. aureus* SH1000, grown on AGBs, were collected in a 10 ml glass jar using sterile forceps. Fixation was performed by covering the samples with 5 ml of 2.5% glutaraldehyde (glutaraldehyde 25%, Carl Roth, Germany) in 0.05 M cacodylate buffer for 1 hour. The fixing solution was stepwise substituted by 0.05 M cacodylate buffer for further preparation, ensuring the beads never dried out completely. The samples were then rinsed three times with 0.05 M cacodylate buffer, each lasting 10 minutes. Osmium tetroxide, whose primary purpose is contrasting, was used as a secondary fixation agent [98]. Two millilitres of 2% osmium tetroxide (ReagentPlus®, 99.8%, Merck, Germany) in distilled water were added to the samples immediately after removal of the buffer and incubated for an additional 30 minutes. Another washing with cacodylate buffer followed as previously described. Dehydration of the fixed colonised AGBs was performed using alcohol at growing concentrations. This was done as a preparation for the critical point drying with carbon dioxide (CO2). The samples were transferred directly into the critical point dryer’s magazine (Foissner vessel, CPD 030, BAL-TEC-Leica, Germany) for better handling. The alcohol series consisted of 30%, 50%, 70%, 90%, and 2x 100% alcohol with a molecular sieve. Starting with the lowest concentration, the Foissner vessel was placed in a beaker filled with alcohol and incubated for 10 minutes. The dehydration process was continued sequentially, using the highest alcohol concentration, including a molecular sieve. The Foissner vessel with the samples was now placed in the cabin of the critical point dryer, and the remaining water and alcohol were entirely replaced by liquid CO2. Liquid CO2 changes into the gaseous phase at 31°C and 74 bar. In this way, the surface structure of a sample can be preserved, which would be damaged in the presence of water during the transition from the liquid to the gaseous state. After drying, the samples were mounted on 12 mm aluminium stubs and coated with gold using a sputtering system (SCD 040, BALZERS, Liechtenstein). For an even gold layer on the preparations, the air in the coater was replaced by inert Argon. The samples were then placed in the SEM and recorded at working distances from 6.5 mm to 12 mm and magnifications between 50 and 27 k-times.

### Confocal fluorescence microscopy

Initially, the AGBs were cultivated with the *E. coli* GFP-expressing strain, as described before. After maturation and a last washing step with saline, the samples were transferred initially into a chambered coverslip with 8 wells and a glass bottom for high-end microscopy (µ-Slide 8 Well, Ibidi, Germany). The beads in the chambers were covered with saline, the coverslip was closed with a lid, and the images were acquired with an inverted confocal laser scanning microscope (SP8-1, Leica, Germany) equipped with a 40x objective. GFP was excited from 488 nm and detected at 510 nm. Both single images and Z-stacks were created from the bead surface or cross sections and later edited in ImageJ.

For live fluorescence microscopy, we follow the same procedure as described before for E. *coli* for confocal microscopy with the three species of GFP-expressing strains with some variations. We have developed a 3D adapter to stabilise the beads on a normal microscope coverslip (60x 25 mm) for microscope observation of biofilms on 4mm AGBs. The adapter was designed in AutoCAD and printed using a Form 4B 3D printer (Formlabs, USA). It has precise external dimensions of 25mm x 25 mm and a central 5 mm hole to hold 4mm beads securely (Fig. S4). The 3D_bead_adapter.stl file is also provided as a supplementary file. This adapter provided a stable and optimal imaging surface for live microscopy of biofilm of *E. coli*, *P. aeruginosa* and *S. aureus*, which was performed employing a DMi 8 inverted microscope (Leica, Germany) equipped with a 100X oil immersion objective (Fig. 3B).

### Coupling of melittin to the agarose-gelatine beads

Using the same procedure that is employed for affinity chromatography of crosslinked agarose media, we modified the method described by March *et al* [99] to couple melittin with the agarose beads. The washed agarose beads, as described in a previous section of this manuscript, were first activated with cyanogen bromide. A volume of 500 µl of coupling buffer (0.1 M Sodium bicarbonate, pH 9.5 in distilled water) along with the amino ligand to be coupled, in our case melittin (5 mg/ml, Genescript, USA) and Bovine serum albumin (BSA) (1 mg/ml, Merck) was used per bead. The coupling reaction was carried out for 20 hours at 4°C. Thereafter, the beads were washed with 20 volumes of buffer containing 0.1 M sodium acetate, pH 4, 2 M urea and 0.1 M sodium bicarbonate, pH 10. The beads were then dialysed against 100 volumes of sterile NaCl 9% solution and stored at 4°C. All operations were made under sterile conditions under a laminar flow with filtered sterilised buffer to prevent bead contamination.

### Staphyloxanthin accumulation. *S. aureus*

SH1000 biofilms growing in AGBs or glass beads were grown in 2 ml of M9 medium with moderate shaking at 37°C for 48 hours with a medium change every 24 hours. The M9 medium (Merck, Germany) used for this experiment was supplemented with 0.4% casamino acids (Carl Roth, Germany), 5 mM MgSO4 (Carl Roth, Germany), and 0.5% glucose as carbon sources (final concentrations). Glucose and magnesium salt were added after autoclaving. Cultures were centrifuged for 1 minute at 10,000 x g, and the supernatant was removed. Each bead was washed twice with 1 ml of NaCl 0.9% solution and added 500 µL of methanol and incubated at 55°C for 30 minutes. Tubes were centrifuged for 2 minutes at 10,000 x g. 300 µL from each tube was added to each well of a polystyrene 96-well plate and then quantified at OD450 on a GloMax Explorer plate reader (Promega, USA). Five biological replicates were performed for each group. For normalisation, all OD450 values were divided by the CFU count of each bead. Normalised production of staphyloxanthin was expressed as a ratio of OD450/CFU, and AGBs and glass beads were compared.

### Quantification of the virulence factors pyocyanin and pyoverdine

To investigate how the virulence of *P. aeruginosa* PAO1 is affected by different substrates, we collected the bacterial supernatant from six biofilm-containing samples on glass or AGBs from all wells of a 24-well plate. The collected supernatant was transferred into 15 ml Falcon tubes and then centrifuged at 4,000 x g for 15 minutes at 20°C. After centrifugation, the resulting supernatant was separated from the bacterial pellets and used for further purification and quantification of pyocyanin concentration. To extract Pyocyanin, we followed a method with some modifications. We incubated 5 ml of the supernatant from each sample with 1 ml chloroform for 10 minutes, with vortexing in between. This process formed an organic phase (blue colour) at the bottom of the tubes. We then pipetted 900 μl from the organic phase and transferred it into 1.5 ml microcentrifuge tubes. After centrifugation for 1 minute at 10,000 x g to remove impurities, 700 µL of the organic phase was transferred to a new microcentrifuge tube and further extracted with 700 µl 0.1N HCl solution. From each sample, 7×100 µl of the aqueous phase formed at the top (red colour) was transferred to a polystyrene 96-well plate. Absorbance was measured at OD520, with the 0.1N HCl solution used as a blank. We performed five experimental replicates for each group. To normalise the results, we divided all measured values by each experiment’s average group CFU counts. The normalised production of pyocyanin was expressed as a ratio of the measured values to CFU and then compared. For pyoverdine, we used an identical setup as for pyocyanin, with identical culture conditions and number of replications. The 24-hour cultures’ supernatants were centrifuged at 10,000 x g for 10 minutes, and the pigment level was determined by measuring its fluorescence, with excitation at 405 nm and emission at 450 nm.

### AHL level determination

The AHL activities of the *P. aeruginosa* PAO1 were assessed using a previously described colourimetric method [74]. Briefly, the bacterial isolates were cultured overnight in 5 mL of sterile Muller-Hinton broth (Merck, Germany) at 37 °C. Then, 1.5 ml of the overnight cultures were aseptically transferred into sterile centrifuge tubes (Eppendorf, Germany) and centrifuged at 10,000 x g for 15 min. The cell pellets were discarded, and this process was repeated three times. The resulting supernatants were filtered through 0.2 μm membrane filters (Sartorius, Germany) to remove any cell debris. The filtrates were mixed with ethyl acetate and shaken for 10 min. Afterwards, the mixture was left to stand for five minutes in a separating funnel, resulting in the formation of an upper organic layer and a lower aqueous layer. The upper layer was collected in sterile tubes, while the lower portion was subjected to two additional extractions following the same procedure. The collected upper portions from each sample were combined and dried in an oven at 40°C. For liquid-liquid extraction (LLE) [100], 40 µL of each dried extract was prepared and inoculated into 96-well polystyrene flat-bottom tissue culture microplate wells. Then, 50 μL of a 1:1 mixture of hydroxylamine (2 M) and NaOH (3.5 M) was added to each well and mixed with the sample. Subsequently, an equal amount of a 1:1 mixture of ferric chloride (10% in 4 M HCl) and 95% ethanol was added. The OD was measured at 520 nm. The AHL extracts were concentrated by drying in an oven at 40°C overnight and stored at −20°C for further analysis. The presence of lactone compounds was indicated by a dark brown colour in all samples, although in some cases, repetitive pipetting of the mixtures caused a colour change to yellow, depending on the lactone concentrations. To ensure that an acidic pH was maintained, we monitored the pH throughout the experiment, as under alkaline conditions, AHLs undergo rapid inactivation through pH-dependent lactonolysis. This reaction hydrolyses the homoserine lactone ring, resulting in the open-ring form corresponding to N-acyl-homoserine. It’s worth noting that this lactonolysis reaction can be reversed by acidification.

### Agarose-gelatine stiffness measurement

The stiffness of agarose-gelatine beads, both with and without crosslinking, was measured using a static compression test described elsewhere [101]. The beads (4 mm) were directly used for the measurement. Then, beads were placed at room temperature overnight for temperature equilibration before the mechanical testing. Young’s modulus was determined by fitting a line to the linear region of the stress-strain curve. Five samples were used for each measurement.

### Sample preparation for proteomics of bacterial biofilms

Each experimental condition consisted of six replicates. A volume of 500 µl of denaturation buffer (6M urea/2 M thiourea in 10 mM HEPES pH 8.0) was added into each sample consisting of three beads of glass, agarose-gelatine and polystyrene substrate from multiwell plates. The samples went for five freeze-thaw cycles to cause lysis and protein release. The resulting lysate was used for in-solution protein digestion as described previously [102]. Briefly, proteins, re-suspended in denaturation buffer, were reduced by the addition of 1 µl of 10 mM DTT dissolved in 50 mM ammonium bicarbonate (ABC) and incubated for 30 minutes, followed by a 20-minute alkylation reaction with 1 µl of 55 mM iodoacetamide. As a first digestion step, Lysyl endopeptidase (LysC, Wako, Japan) resuspended in 50 mM ABC was added to each tube in a ratio of 1 µg per 50 µg of total proteins and incubated for 3 hours. After pre-digestion with LysC, protein samples were diluted four times with 50 mM ABC and subjected to overnight trypsin digestion using 1 µg/reaction of sequencing grade modified trypsin (Promega, USA), also diluted before use in 50 mM ABC. All in-solution protein digestion steps were performed at room temperature. After adding iodoacetamide, the samples were protected from the light until the digestion was stopped by acidification, adding 5% acetonitrile and 0.3% trifluoroacetic acid (final concentrations). The samples were micro-purified and concentrated using the Stage-tip protocol [102], and the eluates were vacuum-dried. For a step-by-step protocol, see Protocol IV of supplementary materials.

### Nano liquid chromatography-mass spectrometry (LC-MS) and data analysis

Peptides were reconstituted in 30 µl of 0.05% trifluoroacetic acid, 4% acetonitrile in water, and 2 µl were analysed by an Ultimate 3000 reversed-phase capillary nano liquid chromatography system connected to a Q Exactive HF mass spectrometer (Thermo Fisher Scientific). Samples were injected and concentrated on a trap column (PepMap100 C18, 3 µm, 100 Å, 75 µm i.d. x 2 cm, Thermo Fisher Scientific) equilibrated with 0.05% TFA in water. After switching the trap column inline, LC separations were performed on a capillary column (Acclaim PepMap100 C18, 2 µm, 100 Å, 75 µm i.d. x 25 cm, Thermo Fisher Scientific) at an eluent flow rate of 300 nl/min. Mobile phase A contained 0.1% formic acid in water, and mobile phase B contained 0.1% formic acid in 80% acetonitrile/20% water. The column was pre-equilibrated with 5% mobile phase B followed by an increase of 5 - 44% mobile phase B in 35 min. Mass spectra were acquired in a data-dependent mode utilising a single MS survey scan (m/z 300–1650) with a resolution of 60,000 and MS/MS scans of the 15 most intense precursor ions with a resolution of 15,000. The dynamic exclusion time was set to 20 seconds, and automatic gain control was set to 3×10^6^ and 1×10^5^ for MS and MS/MS scans, respectively.

MS and MS/MS raw data were analysed using the MaxQuant software package (version 2.0.3.0) with the implemented Andromeda peptide search engine [103]. Data were searched against the reference protein sequence databases downloaded from UniProt: *E. coli* (4448 proteins, Proteome ID UP000000625), *P. aeruginosa* (5564 proteins, Proteome ID UP000002438), and *S. aureus* (2889 proteins, Proteome ID UP000008816). Filtering and statistical analysis were performed using the software Perseus version 1.6.14 [104]. Missing values are imputed using the default settings. Mean log2 fold protein LFQ intensity differences between experimental groups (glass beads vs. AGBs) were calculated in Perseus using student’s t-tests, and *p*-values were adjusted via false discovery rate (FDR) of 0.05 as a cut-off.

## Statistical analysis

The student’s t-test was used for pair comparisons if sample data followed a normal distribution and had homogenous variance. Otherwise, the Mann-Whitney U-test was used instead. For multiple comparisons, we used one-way ANOVA with the Holm-Šídák test for parametric data or the Kruskal-Wallis test for non-parametric ones. All statistical tests were done in GraphPad (Prism, version 9) or R version 4.3.3 [105].

## Acknowledgements

We want to thank the assistance of the Core Facility BioSupraMol (Freie Universität Berlin), which was supported by the Deutsche Forschungsgemeinschaft (DFG).

## Author Contribution

The authors confirm their contribution to the paper as follows: study conception and design: DR and ARR. Data collection and analysis: MH, DR, AN, BK, NP, ARR. Analysis and interpretation of results: MH, DR, AN, BK, NP, JR and ARR. Draft manuscript preparation: DR, MH, AN, ARR. Manuscript editing and correction: all authors. Supervision: JR and ARR. Funding and resources: JR and ARR. All authors reviewed the results and approved the final version of the manuscript.

## Data Availability Statement

The data, including the raw data files are available within the manuscript and the supplementary information files.

## Abbreviations

DVS: divinyl sulphone
AGBs: agarose-gelatine beads
CFU: colony-forming units
SEM: scanning electron microscopy
LC-MS: liquid chromatography mass spectrometry
OD: optical density
GFP: green fluorescent protein

