## Supplementary material for "Crosslinked agarose-gelatine beads as a substrate for investigating biofilms of bacterial pathogens": Suppl_Material.docx

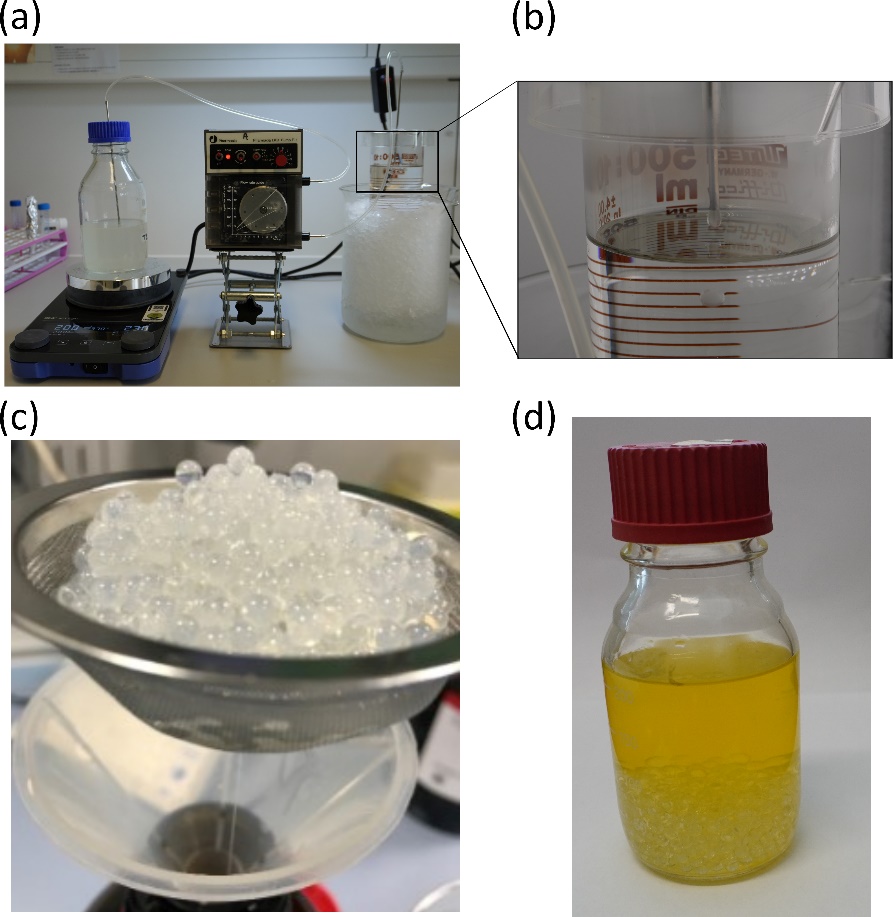

**Fig. S1.** Production process for the agarose-gelatine beads. (A) The bead substrate is carefully placed on a heating plate, with continuous stirring, ensuring the temperature and that the bottle stands higher than the pump and the collection vessel in the ice container. (B) Droplets form at the end of the metal tube following the peristaltic movement rate. The beads gain their spherical form due to hydrogel surface tension, a magnitude drop in temperature, and the inert oil's hydrophobic environment. (C) The resulting beads are sieved and washed with tap water until it flows transparent, with a household detergent added to the first wash to enhance the displacement of oil molecules. The mineral oil is filtered for reuse. The beads are then ready for the following crosslinking steps. (D) The resulting beads, in the medium of choice following autoclaving, can be stored for at least several months at 4°C as necessary. For visualisation, refer also to supplementary video S1.

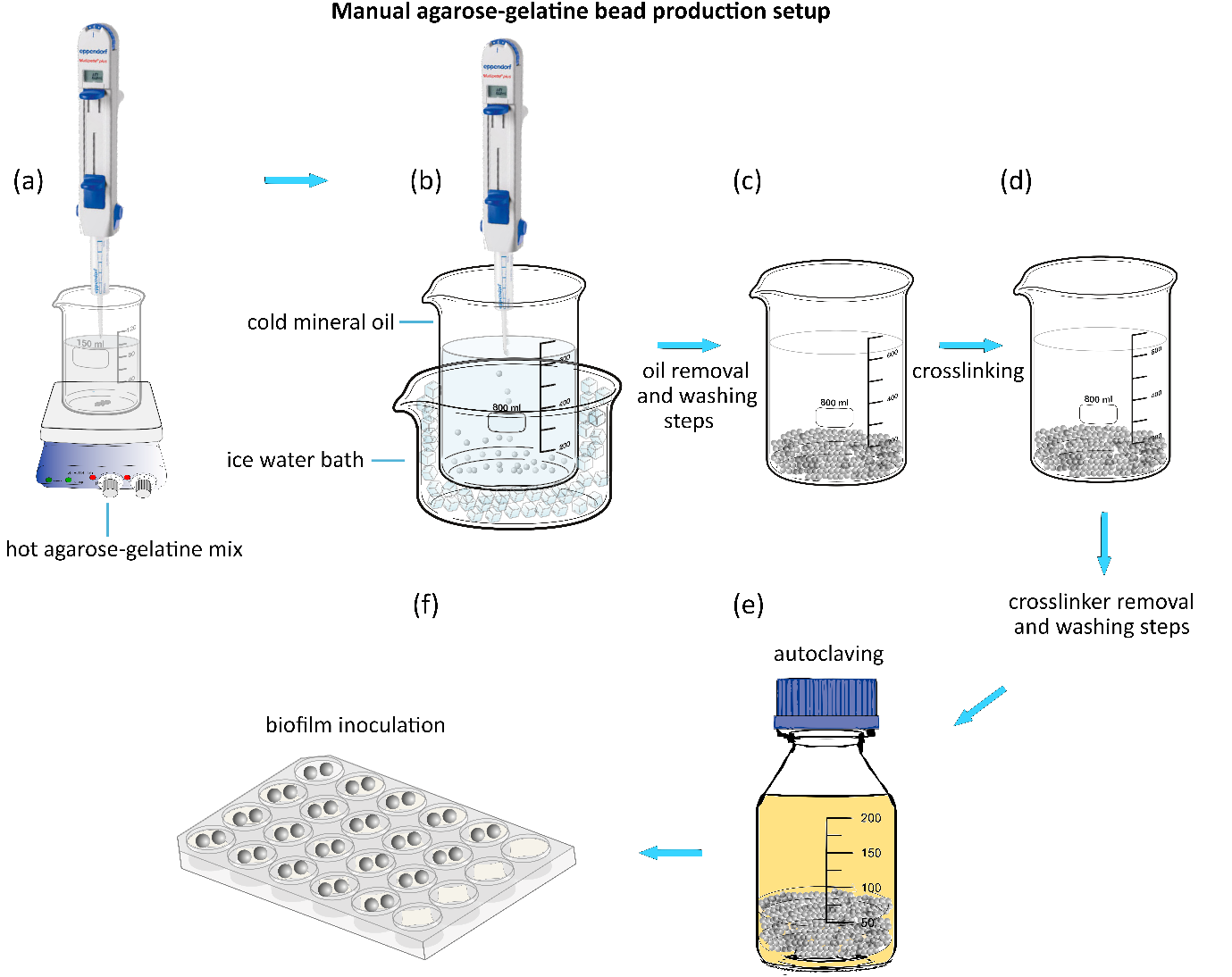

**Fig. S2**. This figure illustrates a more accessible setup for bead production using a multi-step pipette instead of a peristaltic pump, as shown in Fig. 1. In the initial step (a), a fused mixture of gelatine-agarose (2% each in distilled water) is prepared and placed on a magnetic stirrer with a hot plate to maintain liquidity. This solution is continuously added to ice-cold mineral oil using a multi-step pipette to maintain the size and consistency of the beads, followed by the same steps as described in Fig. 1. In step (b), the mineral oil is removed, and washing steps, including the use of detergent, are performed to eliminate oil residues. Following this, in step (c), the beads undergo crosslinking using divinyl sulfone (DVS), with the crosslinker later removed through additional washing steps. After removing the crosslinker, the beads are transferred to a bottle and autoclaved at 121°C for 15 minutes at 1 atmosphere (d), a critical step that ensures the long shelf life of the beads. Once autoclaved, the beads can be stored for several months until use. In the final step (e), the beads are transferred to a 24-well multiwell plate, where the chosen bacteria are inoculated, and the biofilm is established, as previously described. For additional details, see the Materials and Methods section and supplementary video S2.

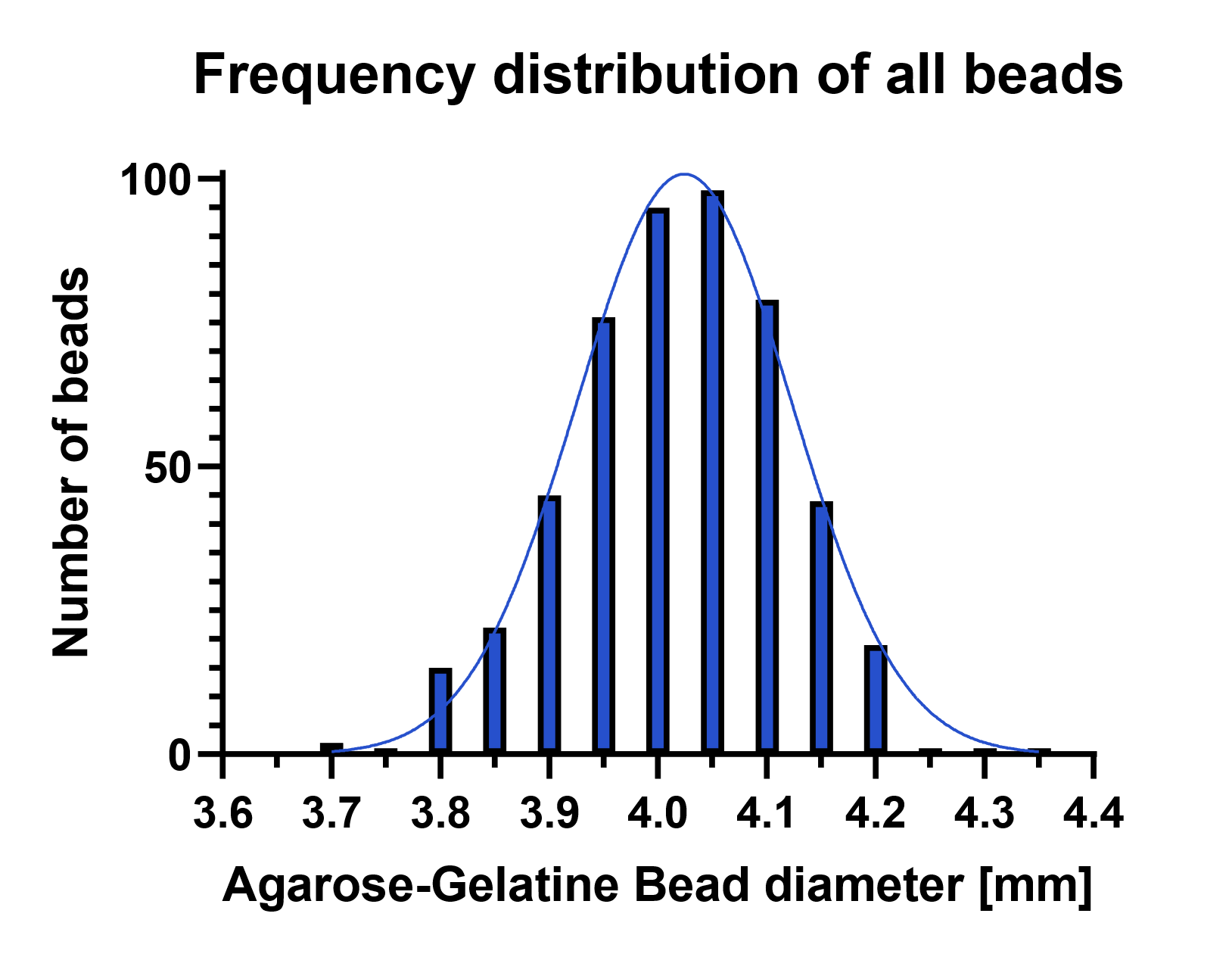

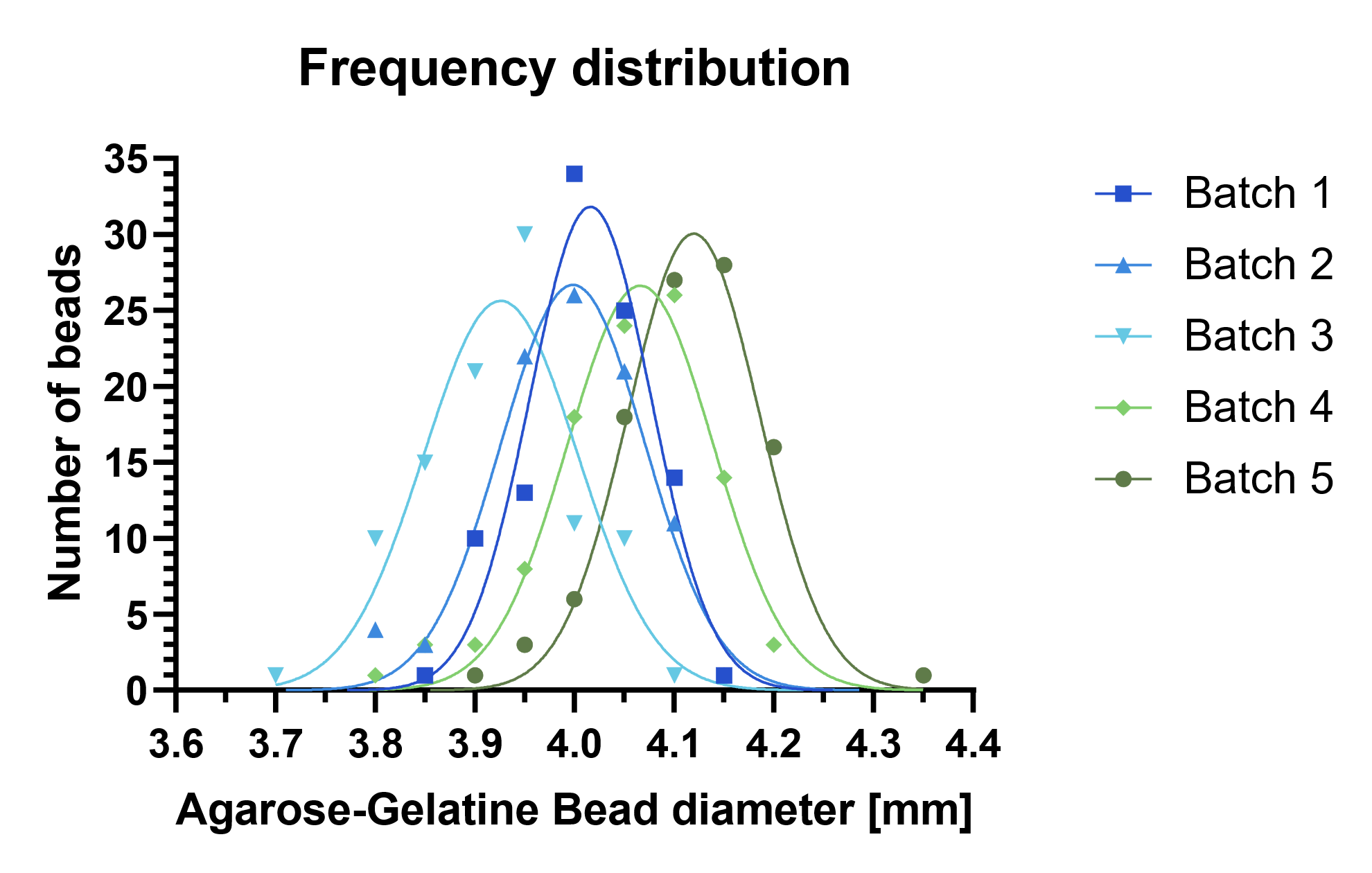

**Fig. S3.** Statistical validation of the consistent size distribution of five independent batches of agarose-gelatine bead random samples (N=100). (A) 99% of the populations of beads in each batch consistently follow a normal distribution as per the D'Agostino & Pearson test with *p*-values above α=0.05. The data plotted here as the number of beads per 0.05 mm for simplification purposes. Means range between 4.11-3.92 mm with standard deviations below 0.08 mm. The frequency distribution of the whole sampled population of agarose-gelatine beads (B) 99% of produced beads follow a normal distribution as represented by the Gaussian curve (B).

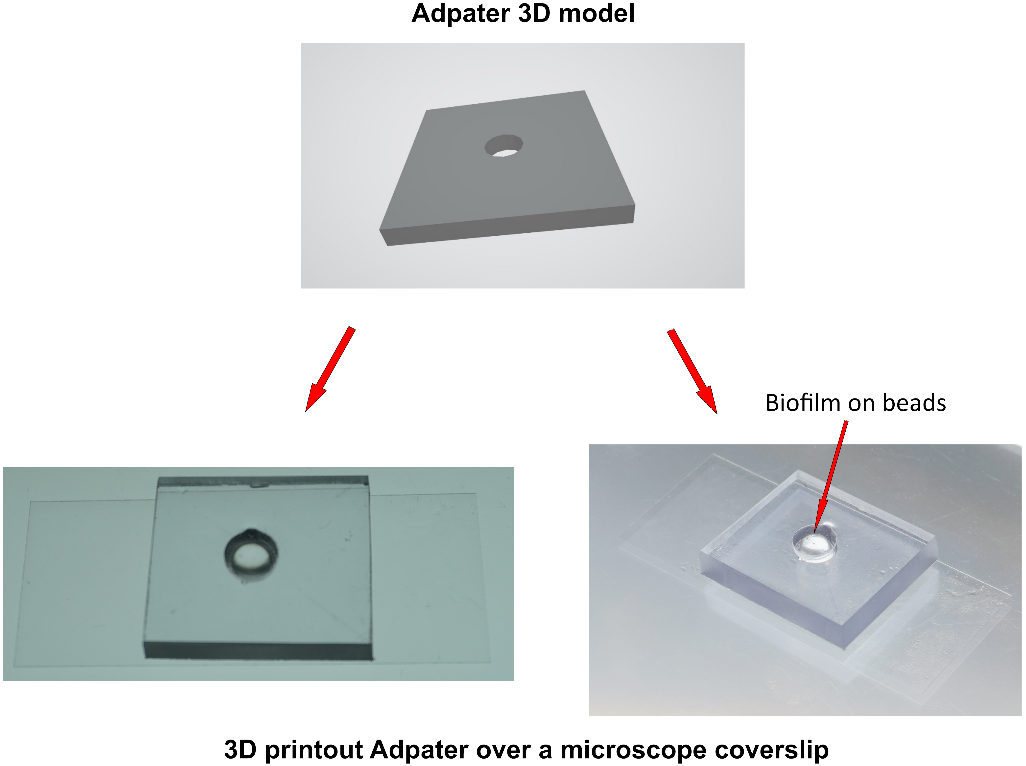

**Fig. S4.** The square 3D-printed adapter (25 mm x 25 mm, 5 mm central hole) was designed in AutoCAD and printed with a 3D printer to create a stable platform for the microscopy of biofilms growing on beads format. Biofilms growing on 4 mm beads were placed in the adapter's central hole, as indicated with a red arrow. A coverslip adheres to the bottom using optical adhesive, providing a flat surface for stable positioning of the specimens. This device is compatible with 100X oil immersion objectives. The 3D_bead_adapter.stl file for microscopy is provided as a supplementary file.

### Supplementary protocol I

**Agarose-gelatine beads production protocol**

**Introduction:** This protocol describes the procedure for producing agarose-gelatine beads (AGBs) as a substrate for various applications in biotechnology and microbiology. Due to their mechanical and thermal stability, AGBs offer a stable matrix for immobilising enzymes, cells, or other biomolecules. The protocol includes steps for bead production, crosslinking to enhance stability, and deactivation to remove residual reactants.

**Materials:**

- Agarose

- Gelatine

- Distilled water

- Inert mineral oil

- Vacuum pump

- Peristaltic pump

- Silicone tubes

- Metal tubes

- 0.5 M sodium carbonate buffer

- Sodium hydroxide

- Divinyl sulphone (DVS)

- 2-Mercaptoethanol

- Glycine

- Sodium chloride

- Parafilm

- Aluminium foil

- Shaker

**Procedure:**

**1. Production of Agarose-gelatine Beads:**

1.1. In a 400 mL bottle, dissolve 4 g each of agarose and gelatine in 200 mL of cold distilled water.

1.2. Autoclave the solution for 20 minutes at 121°C and 2.1 bar absolute pressure.

1.3. Maintain the liquid substrate at 60°C on a hot plate stirrer throughout production.

1.4. Prepare a 1 L glass beaker filled with approximately 500 mL of inert oil as a collection vessel.

1.5. Remove air bubbles from the substrate using a vacuum pump if necessary.

1.6. Cool the collection vessel in a bucket ice water container bath.

1.7. Connect a peristaltic pump and calibrate the flow rate to deliver approximately 40 µl drops, which can be empirically determined using an analytical scale. Perform some adjustments by measuring 10 to 20 beads to reach the desired size.

1.8. Use silicone and metal tubes to deliver uniform drops of the agarose-gelatine mixture into the cooled inert oil. Place the tube approximately 1 to 2 cm from the oil's surface.

1.9. The drops tend to solidify immediately, forming ~4 mm AGBs.

1.10. Sieve and wash the beads with tap water until it runs transparent, filtering and reusing the mineral oil.

**2. Crosslinking of agarose-gelatine beads:**

2.1. Weigh the washed beads to calculate the required amount of DVS (0.5% v/w).

2.2. Cover the beads with approximately 550 mL of 0.5 M sodium carbonate buffer adjusted to pH 11.0 using sodium hydroxide 1 N.

2.3. Add 1.25 mL of the DVS solution per 250 g of wet gel beads in the buffer.

2.4. Allow the crosslinking reaction to proceed overnight on a shaker at room temperature, covering the containing flask with parafilm to prevent evaporation.

2.5. The next day, wash the beads with distilled water by decanting and sieving until they have a neutral pH. Between washes, allow the beads to rest for at least 10 minutes.

**3. Crosslinking reaction stop and crosslinker removal:**

3.1. Mix the wet gel beads with 0.01 mL of 2-Mercaptoethanol per mL of bead suspension in a neutral or slightly alkaline medium.

3.2. Cover the reaction vessel with parafilm to avoid evaporation and aluminium foil to avoid light exposure.

3.3. Allow the reaction to proceed overnight on a shaker at room temperature.

3.4. Discard excess solution and wash beads several times with distilled water.

3.5. Suspend beads in a glycine solution (10% w/v in 1 M sodium carbonate buffer, pH 10.0).

3.6. Allow the reaction to proceed overnight on a shaker at room temperature.

3.7. Wash beads three times with 0.1 M carbonate-bicarbonate buffer (pH 10.0) and then 0.1 M citrate buffer (pH 4.0) with sodium chloride.

3.8. Autoclave beads with media such as LB or Mueller-Hinton broth at 4°C for long-term storage.

**Notes:**

- Beads must remain generally in liquid throughout the production, crosslinking, during and following autoclave and for long-term storage. Desiccation and reconstitution are not possible.

### Supplementary protocol II

**Protocol for cultivating biofilms on agarose-gelatine beads:**

**Introduction:** This protocol provides a method for cultivating biofilms on agarose-gelatine beads (AGBs) for downstream applications such as antibiofilm agent treatments, microscopy, and cell recovery. AGBs provide a stable matrix for biofilm formation and can be used with various bacterial species and growth media.

**Materials:**

- Autoclaved agarose-gelatine beads

- Medium of choice

- 24-well microplate (preferably polypropylene)

- Bacteria culture

- 0.9% NaCl saline solution

- Vacuum aspirator

- Orbital shaker

- 1 ml pipette and tips

**Procedure:**

1. Prepare overnight cultures of the bacteria of choice in the medium of choice.

2. Dilute the overnight culture to achieve an inoculum of approximately 10^6^ colony-forming units (CFU) per millilitre.

3. Add 1 ml of the diluted bacterial culture to each microplate well containing the agarose-gelatine beads and medium.

4. Place two autoclaved agarose-gelatine beads in each well of a 24-well microplate.

5. Place the microplate in an orbital shaker and incubate at 37°C with moderate shaking (20 to 30 r.p.m.).

6. Incubate for 24 to 94 hours, depending on the biofilm's desired maturity and the bacteria's growth speed.

7. If required, perform daily washings to remove planktonic cells and maintain the biofilm by removing the planktonic phase with a 1 ml pipette. Wash the biofilm-containing beads by performing three consecutive washes by gentle addition and removal of with 0.9% NaCl saline solution using a 1 ml pipette. Use a vacuum aspirator for better results.

8. After the desired incubation period, wash the beads as indicated in step 7 and transfer the biofilm-containing beads to a fresh microplate for downstream applications such as antibiofilm agent treatments, microscopy, and cell recovery.

### Supplementary protocol III

**Protocol for extracting viable bacteria from agarose-gelatine beads biofilms.**

**Introduction:** This protocol is a refined method for extracting viable bacteria from biofilm on the surface of agarose-gelatine beads. It includes an additional step of digestion with DNAase I and RNAse A, followed by the addition of papain and mild sonication to digest the extracellular matrix. The protocol allows the determination of anti-biofilm treatment or simplifies the enumeration of bacteria within the biofilm.

**Materials:**

Biofilm-forming bacteria (e.g., *Staphylococcus aureus*, *Escherichia coli*, *Pseudomonas aeruginosa* etc)

Sterile agarose-gelatine beads for biofilm culture

- Digestion buffer (150 mM NaCl, 50 mM Tris-buffer, 10 mM MgCl_2_, 5 mM L-cysteine, 5 mg/ml DNAse I, 2 mg/ml RNAse A, pH 7.5).

- Papain stock solution 10 mg/ml

- 2 ml microcentrifuge tubes

- Microcentrifuge

- Sonicator bath

**Procedure:**

1. Cultivate biofilm-forming bacteria in agarose-gelatine beads according to Supplementary Protocol II under biofilm-inducing conditions until mature biofilms are formed, including those subjected to a chosen treatment (e.g., antimicrobial treatment).
2. Following bead culture, wash the beads with 0.9% NaCl solution.
3. Transfer two beads from each well of the 24-well microplates into separate 2 ml microcentrifuge tubes.
4. Add 200 µl of digestion buffer (150 mM NaCl, 50 mM Tris-buffered saline (TBS) at pH 7.5, 10 mM MgCl_2_, 5 mM L-cysteine, 5 mg/ml DNAse I, and 2 mg/ml RNAse A) to each microcentrifuge tube.
5. Incubate the tubes for 15 minutes at 37°C in a thermo block or recirculating water bath, moderately shaking intermittently to promote thorough enzymatic digestion.
6. After enzymatic digestion, add 20 µl of papain stock solution (10 mg/ml) to each tube, achieving a final concentration of approximately 500 µg/ml. Submerge the tubes in a pre-warmed sonication bath and sonicate for 5 minutes to further disrupt the biofilm matrix.
7. Transfer the supernatant containing the solubilised biofilm components to fresh tubes. Wash the beads once with 1 ml of digestion buffer, then combine the wash with the previously collected supernatant to ensure maximal recovery of biofilm material.
8. Centrifuge the combined suspension at 10,000 × g for 5 minutes to pellet bacterial cells and any remaining debris.
9. Discard the supernatant and carefully resuspend the bacterial pellet in 1 ml of 0.9% NaCl solution.
10. Plate appropriate dilutions of the resuspended bacterial suspension on selective media of choice to determine the colony-forming unit (CFU) count, facilitating subsequent experimental procedures using the recovered bacteria.

### Supplementary protocol IV

**Protocol for preparation of bacterial cell lysate for proteomics (LC-MS) from agarose-gelatine bead biofilms by freeze-thaw cycles.**

**Introduction:**

This protocol describes a method for preparing bacterial cell lysate suitable for proteomic analysis using liquid chromatography-mass spectrometry (LC-MS) from biofilms grown on agarose beads. The protocol involves treating the biofilm-containing agarose beads with desired conditions, followed by freeze-thaw cycles to lyse the cells and extract proteins. The resulting lysate can be used for downstream proteomic analysis to characterise the bacterial proteome.

**Materials:**

- Agarose-gelatin bead biofilm samples

- Dry ice or liquid nitrogen

- Water bath set to 37°C

- Microfuge tubes

- Microfuge

- Vacuum concentrator machine

- HEPES buffer: 1M solution

- 50 mM buffer Ammonium Bicarbonate (ABC). Always use a freshly made solution.

- 55 mM Iodoacetamide solution. The solution should be freshly made and protected from light.

- Dithiothreitol (DTT) 1 M stock solution freshly made.

- Urea buffer (6 M urea, 2 M thiourea, and 10 mM HEPES, pH 8.0).

- Buffer A (5% acetonitrile, 0.3% trifluoroacetic acid). Always use a freshly made solution.

- Buffer B (80% acetonitrile, 0.3% trifluoroacetic acid). Always use a freshly made solution.

**Procedure:**

1. Prepare biofilm-containing agarose beads according to the desired experimental conditions following Supplementary Protocol II. Use at least five replicas per experimental condition, including a control group.
2. Wash the beads with 0.9% NaCl solution following the bead culture.
3. Transfer two beads from each well of the 24-well multiwell plates into separate 1.5 ml microcentrifuge tubes.
4. Add 200 µl of urea buffer (6 M urea, 2 M thiourea, 10 mM HEPES, pH 8.0) to each microcentrifuge tube.
5. Freeze the samples on dry ice or liquid nitrogen and leave for at least 2 minutes. Thaw immediately at 37°C in a nearby water bath. Mix well by flicking the tubes to ensure complete dissolution of urea. Repeat the freeze-thaw process for four more cycles (5 cycles in total).
6. Centrifuge the combined suspension at 10,000 × g for 10 minutes to pellet bacterial debris and agarose beads.
7. Carefully collect 100 µl from each tube without disturbing the pellet, transfer it to a fresh tube, and place it on ice.
8. Use 10 µl of the cell lysate to determine total protein concentration using an appropriate method (Biuret, BCA, or similar).
9. Use approximately 50 µg of total protein for mass spectrometry sample preparation.

**Protein in-solution digestion for mass spectrometry:**

1. Adjust the final volume of the tubes containing the 50 µg of total protein to 50 µl using urea buffer to reach a final protein concentration of 1 mg/ml.
2. Add 2.5 μl of DTT 10 mM solution (see buffers and solutions section) to each tube containing 50 μl of protein solution and incubate for 30 mins at room temperature.
3. Add 2.5 μl of 55 mM iodoacetamide solution to each tube and incubate for 30 min at room temperature in the dark.
4. Dilute samples with 4 volumes of ABC (final urea concentration should be below 2 M) by adding 200 μl of ABC buffer.
5. Add 2 μl of 0.5 μg/μl trypsin (sequencing grade) and incubate overnight at room temperature.
6. Stop trypsin digestion by adding 8 μl of 10% TFA to each sample (the final concentration will be approximately 0.3%) and mixing well by pipetting.
7. Add 14 μl of acetonitrile to each tube (final concentration will be approximately 5%).
8. Proceed with peptide desalting and purification using the StageTip procedure. Digested peptides can also be stored at -20°C until use.

**StageTip for peptide desalting and purification:**

1. Prepare the 200 μl tips (without filter) for peptide binding and micro-purification upon digestion of the protein sample (so-called StageTips). One tip will be used per sample. Label each tip according to the sample code. Using a puncture device such as a biopsy punch to cut and stack two disks inside 200 μl (see Figure 1 for reference).
2. Using an awl, make a hole in the cap of a 2 ml microcentrifuge tube (Eppendorf-type) with a diameter slightly smaller than the broadest part of the pipette tip. Place a tip containing the C18 membrane disks into the 2 ml plastic tube (Eppendorf-type). This tube will be the holder and collecting reservoir for the tips (see Figure 1 for reference).
3. Add 100 μl of methanol (LC-MS grade) to activate each tip.
4. Centrifuge the tip-carrying tubes for 10 seconds at 10,000 g and check that some methanol is retained on the matrix disks. If all solvent goes through, replace the tip with a new one and repeat the procedure. This ensures that the c18 membrane is tightly packed inside the tip.
5. Spin the tip-carrying tubes for 15 seconds at 3,000 x g to remove all methanol.
6. Equilibrate the tips by adding 200 μl of Buffer A and centrifuge at 3,000 g for 5 minutes at room temperature.
7. Spin the tip-carrying tubes at 3,000 g to remove Buffer A. Proceed as soon as possible to the next step to prevent the matrix from drying.
8. Add the acidified samples to their corresponding tips and centrifuge at 3,000 g for 5 minutes at room temperature.
9. Wash the tips by adding 200 μl of Buffer A and centrifuge at 3,000 g for 5 minutes at room temperature.
10. Place the tip into a tip rack and store at +4°C until the desired time for elution and analysis. The samples can be stored for up to three months under these conditions.
11. For sample elution, add 100 μl of Buffer B to each tip and place them in the collection tubes.
12. Evaporate the solvent by vacuum centrifugation and resuspend the peptides in an appropriate volume of the buffer suitable for liquid chromatography-mass spectrometry (LC-MS), e.g., 20 µl of 4% acetonitrile and 0.05% trifluoroacetic acid.

**Notes:**

- Prolonged storage of digested proteins in plastic tubes may result in irreversible loss of peptides due to binding to the tubes' walls.
- Wear gloves during the procedure to avoid sample contamination from the operator’s epithelial cells and to prevent toxicity or irritation caused by some reagents when they come into contact with the skin.
- All solutions used in this protocol should be freshly made before execution as indicated.
- The use of DTT is intended to reduce disulfide bonds and enhance the chance of detecting peptides containing cysteine.
- Iodoacetamide alkylates free thiols of cysteine residues, preventing the re-formation of disulfide bonds. Thus, consider carbamidomethyl on cysteine as a fixed modification during the database search for peptide identification.

### Supplementary protocol V

Protocol for staining and quantifying bacterial biofilm production using a crystal violet stain.

**Materials:**

- Agarose-gelatine bead biofilm samples and controls

- Medium of choice

- 24-well microplate (preferably polypropylene)

- 96-well microplate

- 0.9% NaCl saline solution

- Crystal violet stock solution (from Gram-stain kit)

- 95% Ethanol solution

- Triton X-100 solution

- Orbital shaker

- 1 ml pipette and tips

- Spectrophotometer for absorbance measurement

1. Cultivate biofilm-forming bacteria on agarose-gelatine beads or any other material according to Supplementary Protocol II under biofilm-inducing conditions until mature biofilms are formed, including fresh, uninoculated beads as control. Use some beads without inoculation as a blank control.
2. Following bead culture, gently wash the beads twice with 0.9% NaCl solution.
3. If necessary, transfer to a new 24-well microplate and stain bead biofilm samples with 0.05% Crystal violet in 0.9% NaCl solution, 1 ml per well, for 1 min.
4. Aspire and discard the staining solution using a pipette or a vacuum aspirator.
5. Rinsed the beads several times with preheated (at 45 C) dH2O2 tap water to allow the interspecifically bound crystal violet to diffuse from the internal areas of the beads. Allow the beads to rest in an incubator for 10 minutes until no visible colour is present in the control blank beads. Remove the wash and allow the beads to dry at room temperature for 10 minutes.
6. Solubilize the remaining crystal violet from the beads by adding 1 ml 95% ethanol containing 0.05% (w/v) triton X-100 and incubate at 45°C for 30 minutes on an orbital shaker at room temperature.
7. Transfer 200 μl samples per well of a 96-well microplate and measure optical density at 590 nm using a plate reader or an equivalent machine.

**Notes**:

- Crystal violet dye is toxic and potentially carcinogenic. Use gloves, a lab coat and eye protection.
- A flexible piece of mesh can be used as a sieve during intensive steps, such as washing with tap water, to ensure the beads retain their position and remain in their corresponding well.
- This method with AGB has a relatively high background due to the diffusion of crystal violet dye to the inner part of the bead hydrogel.
